# An empirical approach to developing and testing a traits-based fire ecology framework for bacterial response to wildfires

**DOI:** 10.1101/2022.06.06.495025

**Authors:** Dana B Johnson, Jamie Woolet, Kara M Yedinak, Thea Whitman

**Affiliations:** University of Wisconsin-Madison, Madison, WI; Colorado State University, Fort Collins, CO; Forest Products Laboratory, USDA Forest Service, Madison, WI

## Abstract

Globally, wildfires represent major disturbances, burning millions of hectares annually. Wildfires can restructure soil microbial communities via changes in soil properties and microbial mortality. Fire-induced changes in bacterial communities may influence soil carbon cycling, and recovery to pre-burn community composition and function may take years. We investigated carbon cycling, soil properties, and the importance of three fire-adaptive strategies – fire survival, fast growth, and affinity for post-fire soil environmental conditions – in structuring soil bacterial communities following burns of varying temperatures in boreal forest soils. To identify taxa with each strategy, we simulated burns and incubated soils, tracking respiration and sequencing DNA and rRNA. We then quantified their abundances in the field following wildfires of varying burn severities. The importance of these strategies varies over time and with burn severity. Fire survival has a small but persistent effect on structuring burned soil communities. Fast growing bacteria rapidly colonize the post-fire soil but return to pre-burn relative abundances between one and five years post-fire. Taxa with an affinity for the post-fire environment thrive post-fire, but the effect of this strategy declines by five years post-fire, suggesting that other factors such as vegetation recovery or bacterial dispersal may influence community composition over decadal timescales.

**Graphical Abstract:** 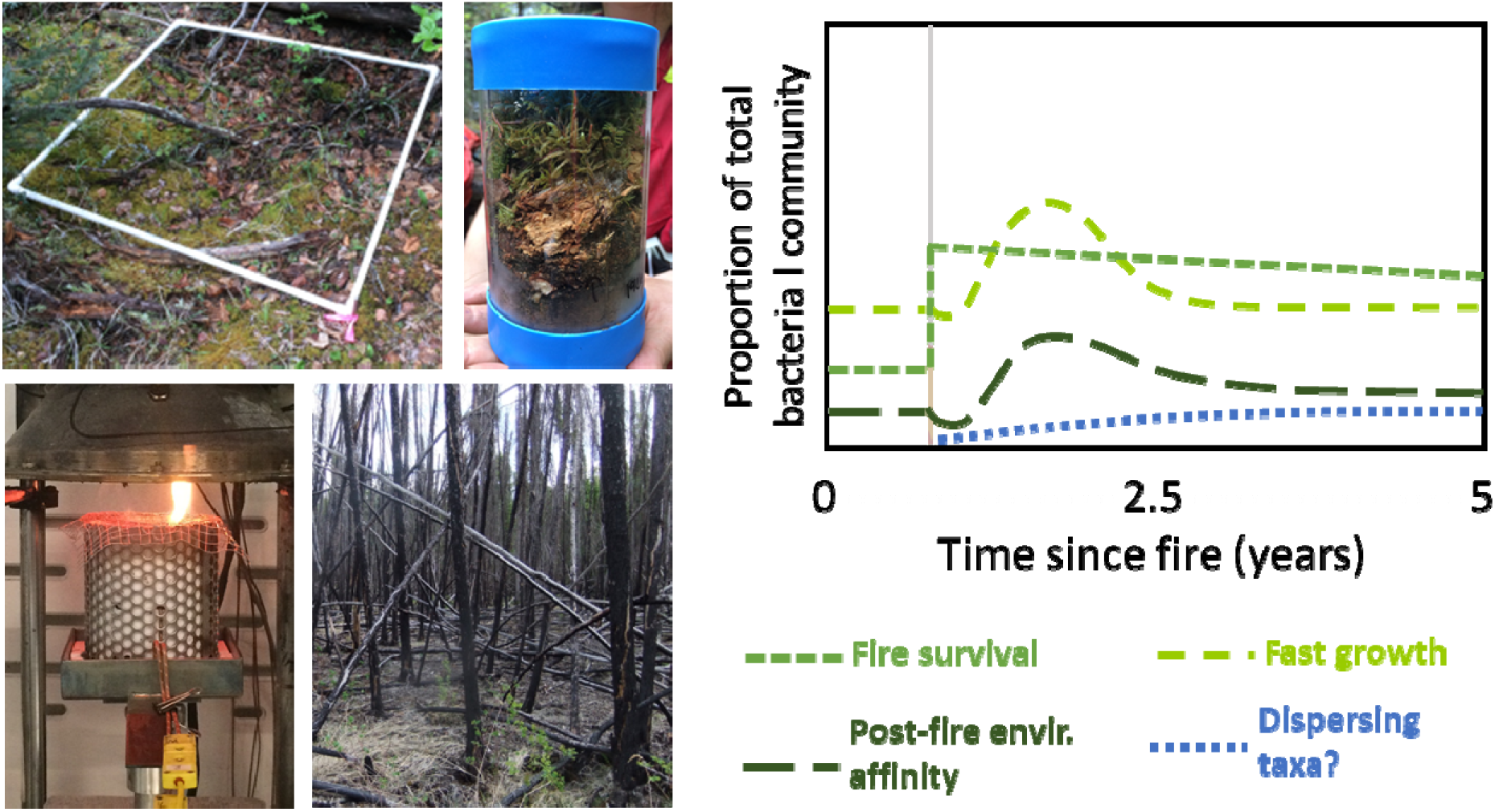

## Introduction

Wildfires are a major ecosystem disturbance of North American boreal forests, burning millions of hectares every year [1]. In this region, wildfire burn severity and area are increasing, driven by climate change [2–4]. During wildfires, belowground soil temperatures can exceed 500 °C, driving carbon (C) and nitrogen (N) loss via volatilization of organic matter (OM) [5–7] and causing a restructuring of soil bacterial communities [8]. Because soil bacteria mediate biogeochemical cycling, the role of fire in restructuring soil bacterial communities is an important component of understanding nutrient cycling and C storage in boreal forest ecosystems. To understand the effects of changing wildfire regimes on soil bacteria, a traits-based framework offers the potential utility of integrating bacteria into biogeochemical models using traits, representing the extremely high diversity of soil bacteria within a few manageable parameters [9, 10].

Building on Grimes’s theories in plant ecology [11], Malik et al. proposed a trait-based framework for soil microbial ecology based on three microbial life strategies: high yield (Y), resource acquisition (A), and stress tolerance (S) [12]. A modified version of this framework may be helpful in untangling the role of fire in structuring soil microbial communities in boreal forest ecosystems adapted to and dependent upon wildfire. We propose three strategies – fire survival, fast growth, and affinity for the post-fire environment – that may confer a post-fire advantage to individual bacterial taxa and roughly map to S, Y, and A strategies, respectively.

Disturbances can be classified as pulses or presses, based on the period of disturbance [13]. Pulse disturbances are characterized by intense yet short-lived environmental pressures. Press disturbances can arise rapidly but persist over a longer period of time. The immediate effects of heat and combustion during wildfires can be considered a pulse disturbance, while longer-term changes to vegetation communities and soil properties, such as pH and C availability [14–16], may be classified as press disturbances for soil microbial communities.

During pulse disturbances of wildfires, soil bacterial survival is affected by temperature and duration of heating, generally decreasing with both [17–20]. Dormancy may be an important trait for maintaining bacterial “seed banks” [21, 22]: soil bacterial mortality can occur across a wide range of temperatures (<80-400 °C) and durations (2-30 min) [17], while dormant cells have been shown to survive temperatures greater than 100 °C for 24 hours in dry conditions [23].

The press disturbance of wildfires on soil bacterial communities lingers long after the flames have receded and soil temperatures return to normal levels. Heat-induced bacterial mortality may open a niche for fast-growing taxa to re-colonize. This is supported by previous work finding a decrease in 16S rRNA copy number months and years post-burn, a parameter that correlates with faster sporulation and growth rates [26, 27]. Bacteria with higher rRNA gene copy numbers may have a selective advantage in responding to increased nutrient availability [25].

Burn-induced changes in soil physical and chemical properties may also advantage bacteria adapted to post-fire conditions. For example, soil pH, which has a strong influence on soil bacterial community composition [28], often increases with burning, and elevated pH has been reported 4-9 years post-fire [14, 29, 30]. Additionally, pyrogenic organic matter (PyOM), produced via the incomplete combustion of soil organic C [31], contains a high proportion of aromatic compounds [32], conferring increased chemical recalcitrance and, thus, persistence [33]. The addition of PyOM to soil can affect microbial community composition [34, 35], although microbial community response is not consistent across varying soil types and with PyOM derived from different sources and production temperatures [36]. Increased pH and/or PyOM, along with other post-fire physicochemical changes, may select for bacteria with an affinity for the post-fire environment.

Our objectives for this study were to assess the relative importance of three fire response strategies – fire survival, fast growth, and an affinity for the post-fire soil environment – in structuring the post-fire soil bacterial community over time. We hypothesized that hours to days after burning, fire survival would be most important in structuring bacterial communities and decrease with time since fire. The importance of fast growth in structuring bacterial communities would supersede fire survival in the weeks to months following burns as fast-growing bacteria recolonize the soil. In the months to years following fire, an affinity for the post-fire soil environment would emerge as the most important strategy structuring bacterial communities. We hypothesized that each of these strategies would have greater effects on bacterial community composition following burns of higher temperature or higher severity [37]. We used simulated burns, soil incubations, and DNA and rRNA sequencing, paired with field data one and five years post-wildfire, to assess advantageous microbial traits for the post-fire environment over time.

## Materials and Methods

For further details on methods, please see Supplementary Information.

### Study overview, site description, and sample collection

For the simulated burns, soil cores were collected from nineteen sites across Wood Buffalo National Park (WBNP), Canada in June 2019 (Fig. 1; Supplementary, Fig. S1, Table S1). The park has both sandy, acidic soils (Eutric Gleysols and Eutric Cambisols, FAO) and organic-rich peatlands (Dystric Histosols, FAO) underlain by discontinuous permafrost [38–40]. The dominant vegetation of the sampled regions of the park is jack pine (*Pinus banksiana* Lamb.), trembling aspen (*Populus tremuloides* Michx.), black spruce (*Picea mariana* (Mill.)), and white spruce (*Picea glauca* (Moench) Voss).

**Fig. 1.**
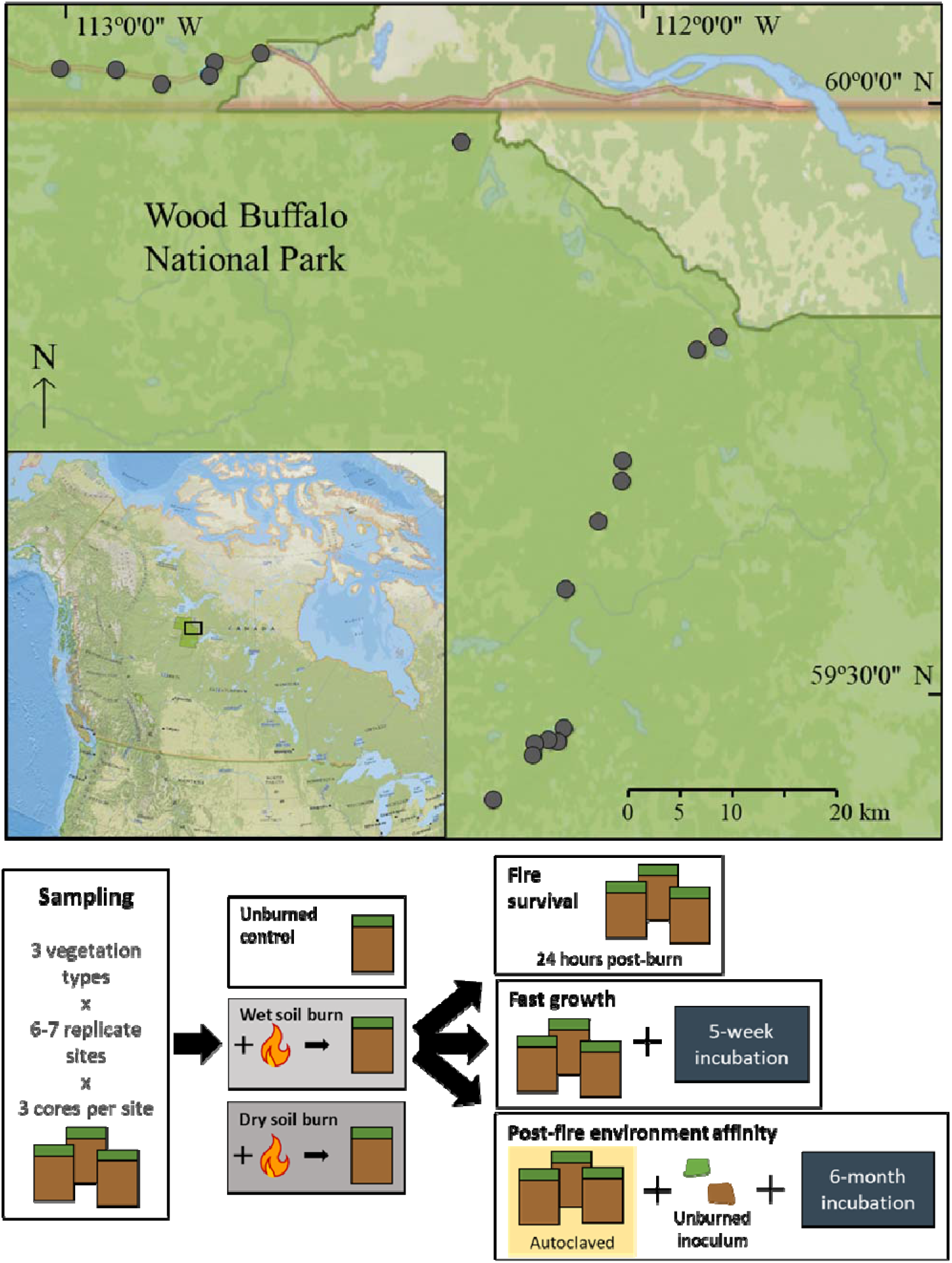
Sampling locations within Wood Buffalo National Park, Canada (top figure). Inset indicates the relative location of the sampling region within North America. Simplified experimental design (bottom figure).

Sites that had not burned in at least the previous 30 years were selected using stratified random sampling using the Canadian National Fire Database (2019), dominant vegetation from the Canadian National Forest Inventory, and soil type from FAO soil survey data [42, 43]. At each site, ten soil cores (15.2 cm x 7.6 cm dia.) were collected across a 2 x 2 m grid. Cores were stored at room temperature or cooler and transported to Madison, WI within 14 days of collection. Cores were then air dried for six weeks to simulate extreme dry conditions with no precipitation [44].

### Fire simulations

For each site, three cores with similar soil horizon thicknesses were chosen for burn simulations. The first core (“wet soil”) was saturated and allowed to free-drain for 24 hours before the burn, while the second core (“dry soil”) was not wet up before the burn. The third core was not wet up and not burned (“control”). Temperatures during and up to 12 hours after the burn were measured with beaded 20-gauge Type K thermocouples (GG-K-20-SLE, Omega, CT, USA) 1 cm above the base of the core and at the transition between the O horizon and top mineral horizon or, for completely organic cores, 5 cm above the core base. Cores were exposed to a cumulative 7.2 MJ m^-2^ radiant heat flux, typical of a mid- to high-range crown fire in this region [45, 46] – 60 kW m^-2^ for 2 minutes in a mass loss calorimeter (Fire Testing Technology Limited, West Sussex, UK) and allowed to cool overnight (Supplementary, Fig. S1).

### Soil property analyses

Twenty-four hours after the burn, mass loss and moisture of each core were recorded (Supplementary, Table S2). Sub-samples of the O horizon and mineral soil were immediately preserved for pH measurements [47] and C and N analysis (Flash EA 1112 CN Automatic Elemental Analyzer (Thermo Finnigan, Milan, Italy)). Soil texture was determined using a physical analysis hydrometer at the UW-Madison Soil and Forage Lab.

### Soil incubations and gas flux tracing

Fire survival was assessed immediately post-burn, as described below. To test for fast growth and for affinity for the post-fire environment, we used two incubation experiments (Fig. 1). For fast growth, we assessed changes in community composition after incubating soils for 5 weeks. For post-fire environment affinity, we wanted to determine which bacteria were suited to the post-fire soil conditions, regardless of their ability to survive the fire, so we inoculated burned soils with microbes from unburned soils from the same site and incubated for six months. Prior to inoculation with 10% dry sample mass basis by horizon [48, 49], we autoclaved soils to reduce total populations. To quantify any artifacts from autoclaving, control incubations were also included using soil from each core that was inoculated without having been autoclaved.

For both incubations, mineral soil from each sample (where present) was packed into a 60 mL glass jar and then topped with soil from the sample’s O horizon to recreate original soil profiles proportionally, packing them to the bulk density of the original core, and maintaining at optimal soil moisture conditions. CO_2_ efflux was tracked over the course of the incubations using KOH traps [50].

### Nucleic acid extraction and sequencing

For fire survival, RNA and DNA were extracted from soil collected 24 hours post-burn using the RNeasy PowerSoil Total RNA Kits and RNeasy PowerSoil DNA Elution Kits (QIAGEN, Germantown, MD, USA) following the manufacturer’s instructions. Residual DNA contamination was removed from the RNA extracts using DNase Max Kits (QIAGEN, Germantown, MD, USA) following manufacturer’s instructions. RNA reverse transcription was carried out using Invitrogen SuperScript IV VILO Master Mix (ThermoFisher Scientific, Waltham, MA, USA) following manufacturer’s instructions. For fast growth and post-fire affinity incubations, DNA was extracted from soil collected at the end of the incubations using the DNeasy PowerLyzer PowerSoil DNA Extraction Kit following the manufacturer’s instructions.

Copy DNA and DNA were amplified via triplicate PCR, targeting the 16S RNA gene v4 region with 515f and 806r primers [51] with barcodes and Illumina sequencing adapters added as per Kozich et al. (2013). PCR amplicon triplicates were pooled, purified, normalized, and sequenced using 2×250 paired end Illumina MiSeq sequencing at the UW-Madison Biotechnology Center (Madison, WI, USA).

### Sequence processing and taxonomic assignments

We quality filtered, trimmed, dereplicated, learned errors, picked operational taxonomic units (OTUs), and removed chimeras using dada2 [53] as implemented in QIIME2 [54]. Taxonomy was assigned using the naïve Bayes classifier [55] in QIIME2 with the aligned 515f-806r region of the 99% OTUs from the SILVA database (SIVLA 138 SSU) [56–58].

### Statistical analyses and trait assignments

We worked primarily in R [59] relying extensively on R packages *phyloseq* [60], *dplyr* [61], *ggplot2* [62], and *vegan* [63]. We identified taxa with the strategy of interest (survival, fast growth, or post-fire environment affinity) from each experiment using the *corncob* package [64] in R. We estimated the log_2_-fold change in the relative abundance of bacterial taxa enriched in burned vs. unburned soil 24 hours post-burn (designated “fire surviving”), taxa enriched at 5 weeks versus 24 hours (designated “fast growing”), and taxa enriched in burned vs. unburned soil following inoculation and a 6 month incubation (categorized as having a post-fire environment affinity). We used a false discovery rate cutoff of 0.05 to adjust p values of differentially abundant taxa to account for false positives. We excluded burned soil cores that did not reach temperatures > 50 °C [17].

For CO_2_ flux data, we fit two-pool exponential decay models for each sample, estimating respiration rate constants and fractional sizes of fast and slow C pools [65] (Eq. 1),

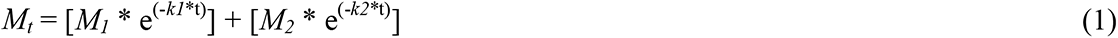

where *M_t_* is the total C pool, *M_1_* and *M_2_* are the fast and slow C pools, respectively, *k1* and *k2* are the respiration rate constants for the fast and slow C pools, respectively, and *t* is time. This model was fit using the *nls.lm* function in the *minpack.lm* package [66] in R.

To compare soil properties across horizons and burn treatments, we used ANOVA, Wilcoxon rank sum tests, and fitted linear models using the *stats* package [67] in R. To assess predictive factors for whole-community composition, we used PERMANOVA on weighted UniFrac distances [68] calculated from a phylogenetic tree built with FastTree [69] from sequences aligned using the *mafft* program [70, 71].

We used the rrnDB RDP Classifier tool (version 2.12) to obtain a mean 16S rRNA gene copy number for each genus in the fast growth dataset [72] and calculated community weighted mean rRNA gene copy number (see Supplementary for details) [25].

### Quantifying post-fire strategies in field data

We quantified the total relative abundance of taxa assigned to each of the strategies using our laboratory experiments in a dataset of natural wildfires in the same region [8, 73]. Briefly, soils were collected from organic and mineral horizons in the field, one year and five years post-fire, across a range of burn severities including unburned sites, and spanning the same vegetation communities and soil types from which cores were collected for the laboratory experiments. OTUs were identified for the field data using similar methods as for the laboratory experiments (same primers, library preparation, sequencing pipeline, and OTU-picking algorithms) [51–53, 56, 58, 74, 75], allowing us to merge the datasets using exact matches for OTUs (amplicon sequence variants (ASVs)). To determine whether there was a significant correlation between burn severity and the total relative abundance of all taxa with a given strategy, we fit linear models, including interaction terms between years since fire (one or five) and burn severity index [76]. If the interaction was not significant, we report results from the model without the interaction.

## Results

### Impact of burning on soil properties

In the dry soil burns, fire propagated downwards from the soil surface as indicated by maximum thermocouple temperatures reaching upwards of 600 °C at the organic-mineral interface (Fig. 2A; Supplementary, Table S3). The thermocouple temperatures in the wet soil burns were much lower (max. recorded=42 °C). Soil pH was significantly higher after dry soil burns in organic horizons (mean=7.2 ± 1.0) than in the unburned control organic horizons (mean=4.5 ± 0.7) (ANOVA, p << 0.001) (Fig. 2B; Supplementary, Table S4). C:N ratios were lower in organic horizons after dry soil burns (mean=12.6 ± 4.3) than in the unburned (mean=20.6 ± 5.9; ANOVA, p << 0.001) or wet soil burns (mean=19.9 ± 4.7; ANOVA, p << 0.001) (Fig. 2B; Supplementary, Table S4). Decreased C:N was largely driven by decreased total C content – organic horizons subjected to the dry soil burn had a mean decrease in C of 11.7% (ANOVA, p=0.015) compared to unburned soil, whereas mean total N content was not significantly affected by burn treatment in either organic or mineral horizons (Supplementary, Fig. S2).

**Fig. 2.**
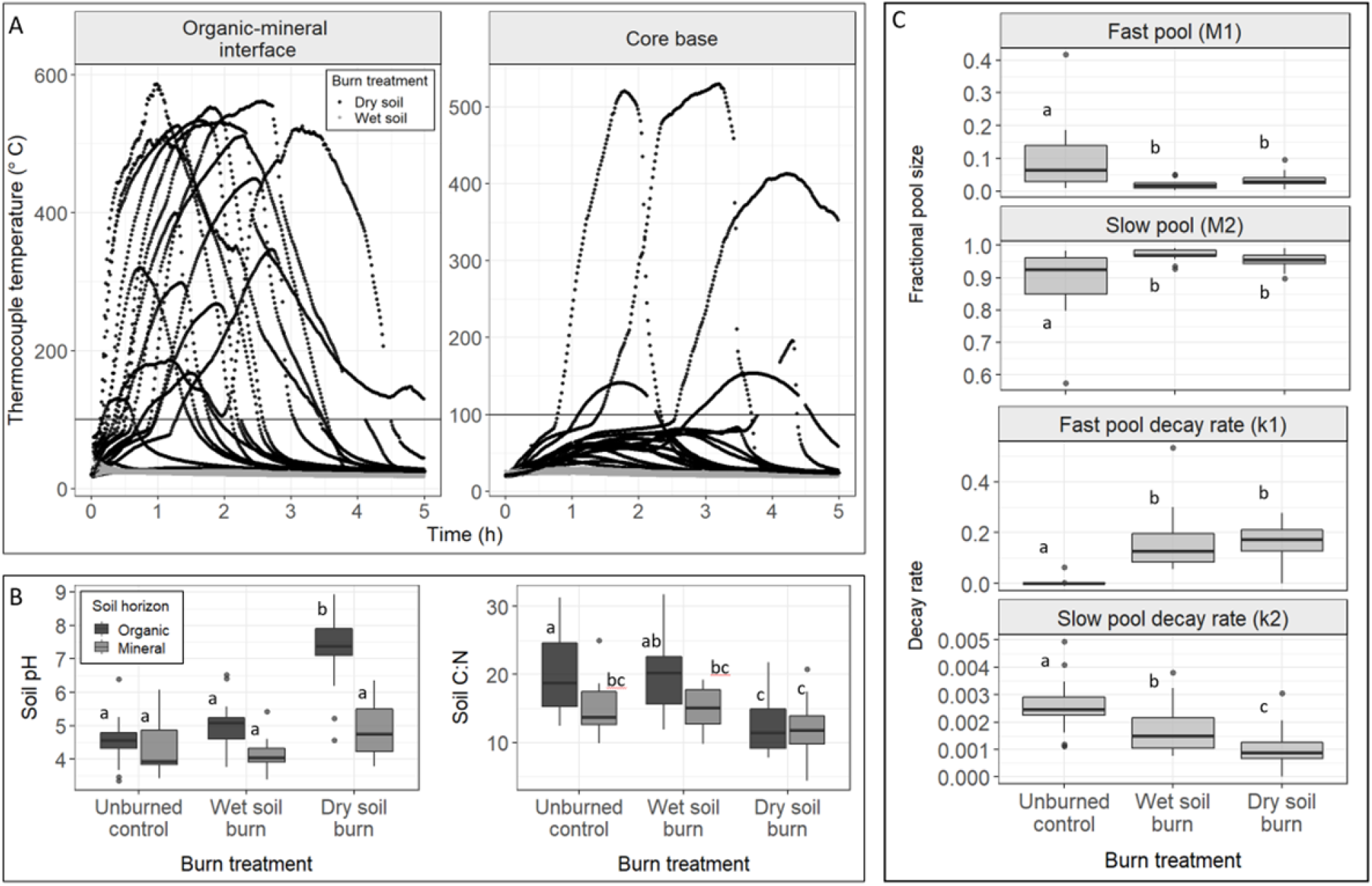
(A) Thermocouple temperatures at the organic-mineral soil interface (left) and 1 cm above the core base (right) during the 5 hours following the initiation of the burn treatments. Burn simulations were initiated at time 0. Temperatures greater than 100 °C suggest the presence of flaming combustion and are not a quantitative measure of soil matrix. (B) Soil pH (left) and C:N (right) across the three burn treatments. Dark fill indicates organic horizon samples while light grey indicates mineral horizons. Different letters indicate significant differences across all treatments. (C) Coefficients for two-pool exponential decay models fit to soil C respiration data (R^2^ of models 0.98-1). Fractional size of the fast (*M_1_*) and slow (*M_2_*) C pool (top two panels) and decay rate coefficients for the fast (*k1*) and slow (*k2*) C pool (bottom two panels) across burn treatments for the 5-week fast-growth incubation. Different letters indicate significant differences across treatments.

### Soil microbial respiration

Two-pool exponential decay models resulted in good fits to respiration data for all samples (R^2^ of 0.98 to 1) (Supplementary, Fig. S3 & S4, Table S5 & S6). In the fast growth incubation, the fractional size of the fast C pool (*M_1_*) was significantly smaller in the burned cores (mean=0.027 ± 0.019; (Wilcoxon signed-rank test, p < 0.001)) than in the unburned cores (mean=0.096 ± 0.10) (Fig. 2C). The fast C pool decay rate coefficient (*k1*) was higher in the burned cores (mean=0.16 ± 0.096) than in the unburned cores (mean=0.0036 ± 0.015; Wilcoxon signed-rank test, p < 0.001). C flux results for the post-fire environment affinity incubation were similar (Supplementary, Table S5 & S6, Fig. S5).

### Community composition in laboratory experiments and the field dataset

There was good correspondence in overall community composition between our lab samples and our field samples (Fig. 3).

**Fig. 3.**
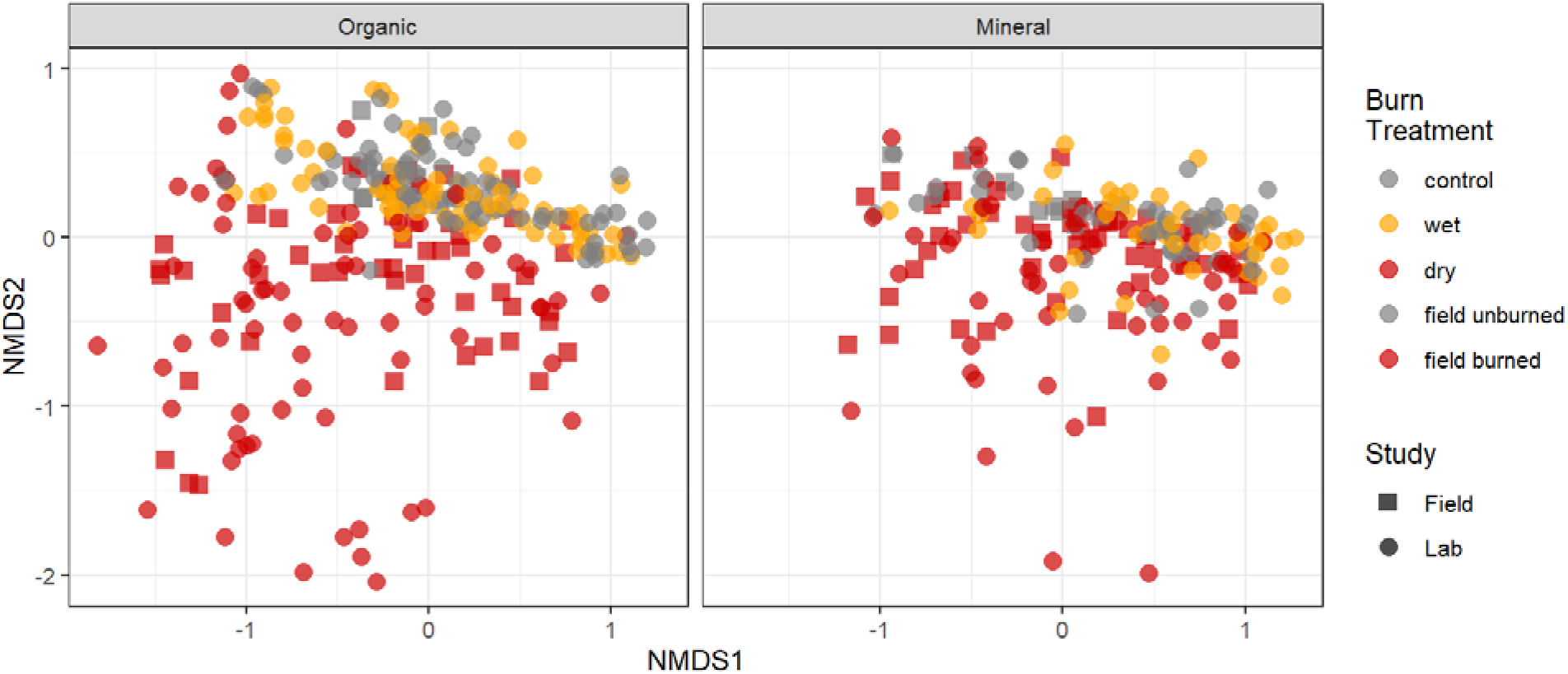
First two axes of NMDS ordination of weighted UniFrac dissimilarities for bacterial communities for mineral (left panel) and organic (right panel) horizons from lab study samples (24-hour incubation, DNA only; five-week incubation; autoclaved post-fire affinity incubation) and field samples (one and five years post-fire) (k = 3, stress = 0.13). Grey shapes indicate unburned lab or field samples, red shapes indicate dry soil burn or field-burned samples, and yellow shapes indicate wet soil burn samples. Circles indicate samples from the lab experiments and squares indicate samples from the field datasets.

For all communities across the three laboratory experiments, all tested factors (dominant vegetation, soil horizon, pre-burn horizon thickness, pH, total C and N, soil texture, and burn treatment) were significant predictors of community composition in combined models (PERMANOVA, p < 0.05 for all factors; Supplementary, Fig. S6, Tables S7-10).

### Fire survival in the lab and in the field

Burning strongly decreased total RNA (10X) and DNA (7X) concentrations for dry soil burns in the organic horizon, and decreased DNA concentrations (3X) in the mineral horizon (Fig. 4A; p ≤ 0.0001).

**Fig. 4.**
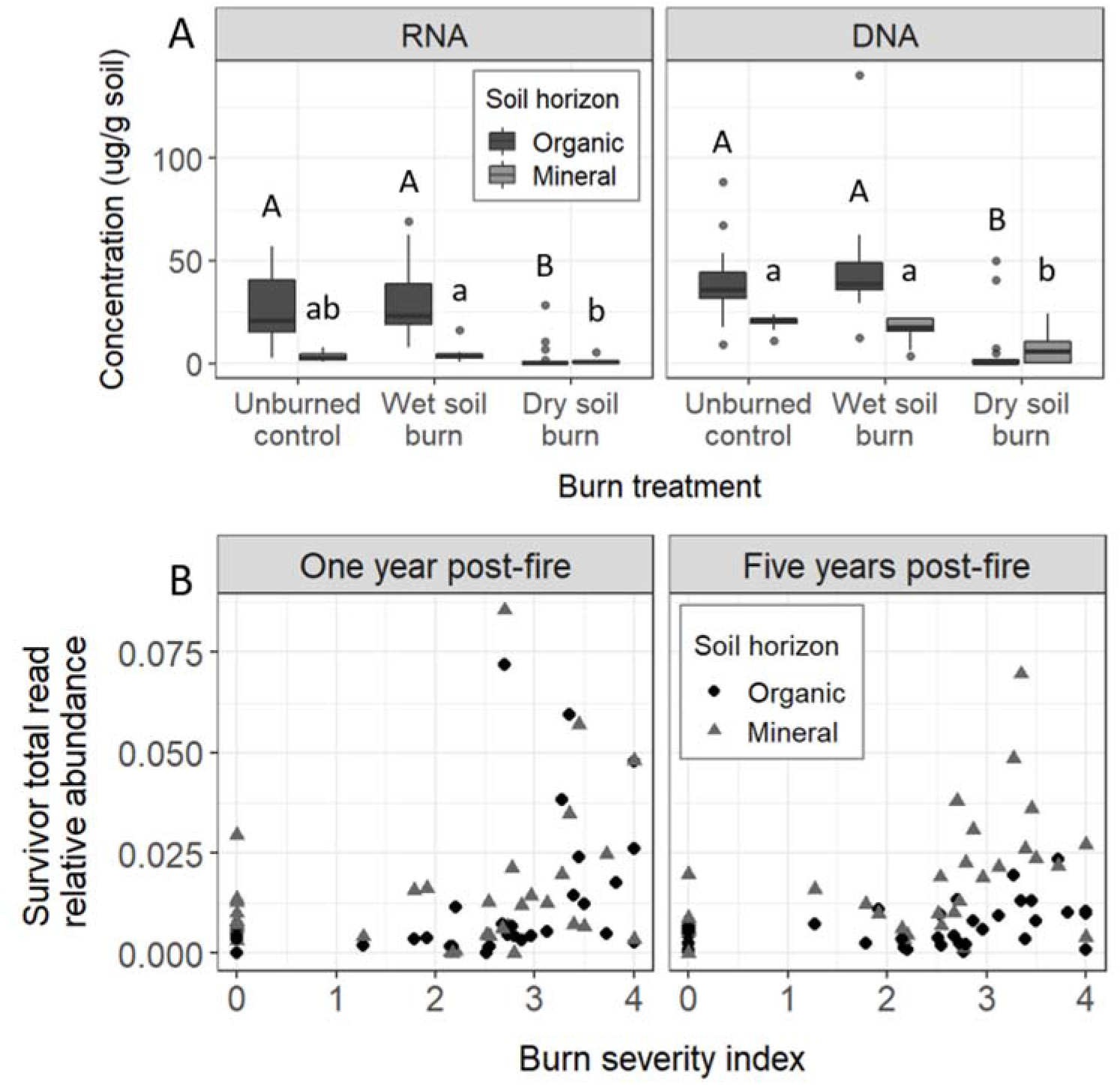
(A) Concentration of total RNA (left panel) and DNA (right panel) extracted from soil 24 hours post-burn. Dark and light grey indicate organic and mineral horizon soil, respectively. Different letters indicate significant differences between treatments for each horizon. (B) Total relative abundance of lab-identified fire-survivor taxa in field data one (left) and five (right) years post-fire, *vs*. burn severity index. Black circles indicate organic horizon samples while grey triangles indicate mineral horizon samples.

17 bacterial OTUs were identified as significantly enriched (“fire surviving”) in soil cores that experienced temperatures > 50 °C compared to unburned control cores (Supplementary, Table S11). These fire-survivor taxa made up a maximum of 9% of the reads in burned communities from the field and were significantly positively correlated with burn severity index (p=0.0002, R^2^ _adj_=0.09), both one and five years post-fire (interaction term between years post-fire and burn severity index p=0.71) (Fig. 4B).

### Fast growth in the lab and in the field

Differences in weighted mean predicted rRNA gene copy number emerged after 5 weeks of incubation: dry soil burns had significantly higher weighted mean predicted rRNA gene copy number (mean=3.4 ± 0.9, organic; mean=2.7 ± 0.9, mineral) than unburned soil (mean=2.0 ± 0.4, organic; mean=1.7 ± 0.1, mineral; ANOVA, p_adj_ < 0.001; Fig. 5A).

**Fig. 5.**
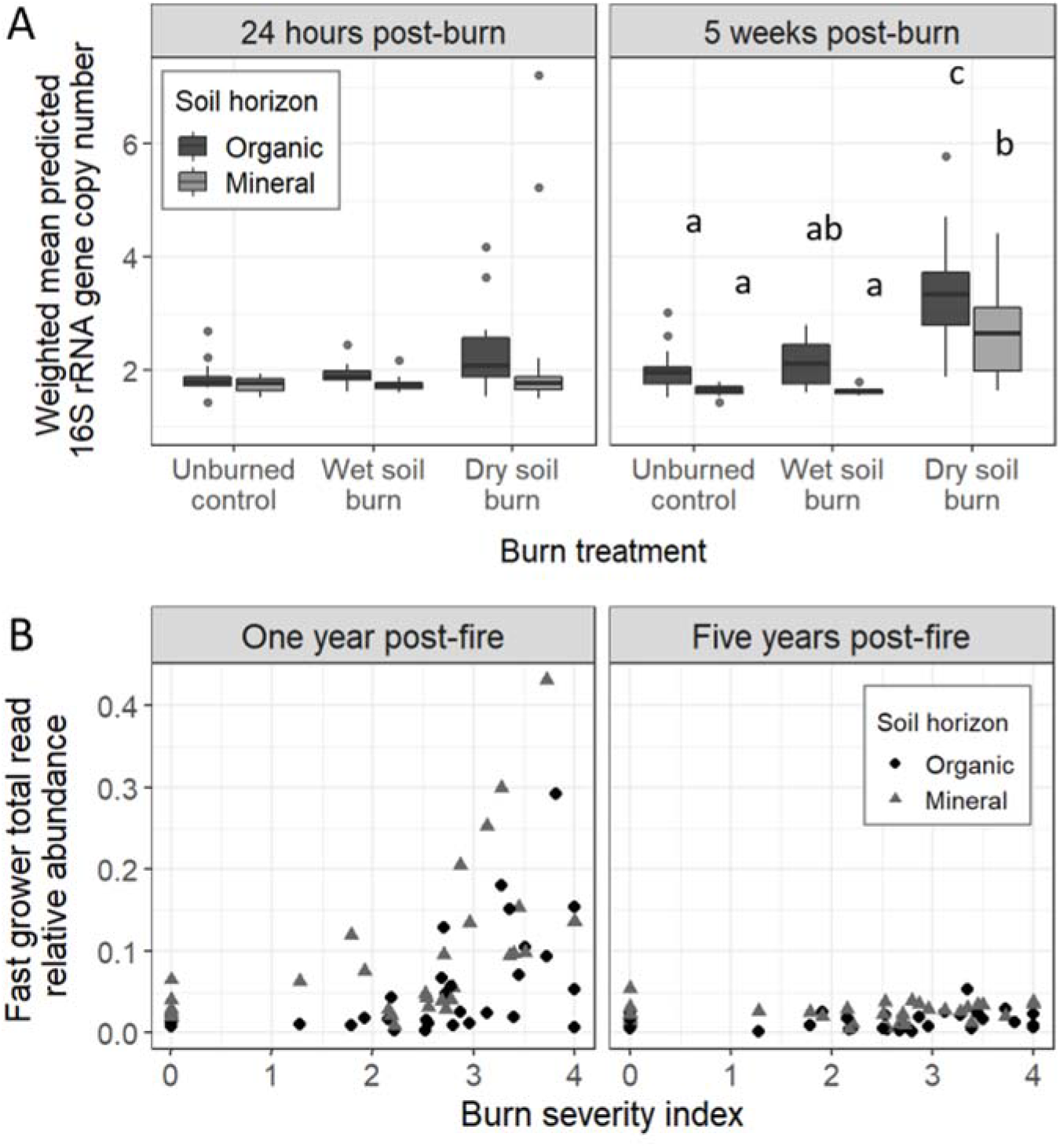
(A) Weighted mean predicted 16S rRNA gene copy number for bacterial communities in the organic (dark grey) and mineral (light grey) soil 24 hours (left) and 5 weeks (right) post-burn for different burn treatments. Different letters indicate significant differences across burn treatments and soil horizons. (B) Total relative abundance of lab-identified fast-grower taxa in field data one (left) and five (right) years post-fire, *vs*. burn severity index. Black circles indicate organic horizon samples while grey triangles indicate mineral horizon samples.

488 OTUs were identified as enriched 5 weeks (versus 24 h) post-burn (designated “fast growers”) (Supplementary, Table S11). Of these OTUs, 9 were also identified as fire surviving. These fast-growing taxa made up a maximum of 43% of the reads in burned communities from the field and were significantly positively correlated with burn severity index (p < 0.001, R^2^_adj_=0.34; Fig. 5B). However, this relationship was completely gone five years post-fire (interaction term between years post-fire and burn severity index p < 0.001).

### Post-fire environment affinity in the lab and in the field

In organic horizons, community compositions changed more with burning at sites where pH increased more after dry soil burns (R^2^ =0.5; p=0.0008; Fig. 6A). The autoclaving treatment (vs. inoculated but not autoclaved) provided the least explanatory power for community composition (R^2^=0.02; Supplementary, Table S10), suggesting soil properties played a larger role in structuring communities than any artifacts of autoclaving.

**Fig. 6.**
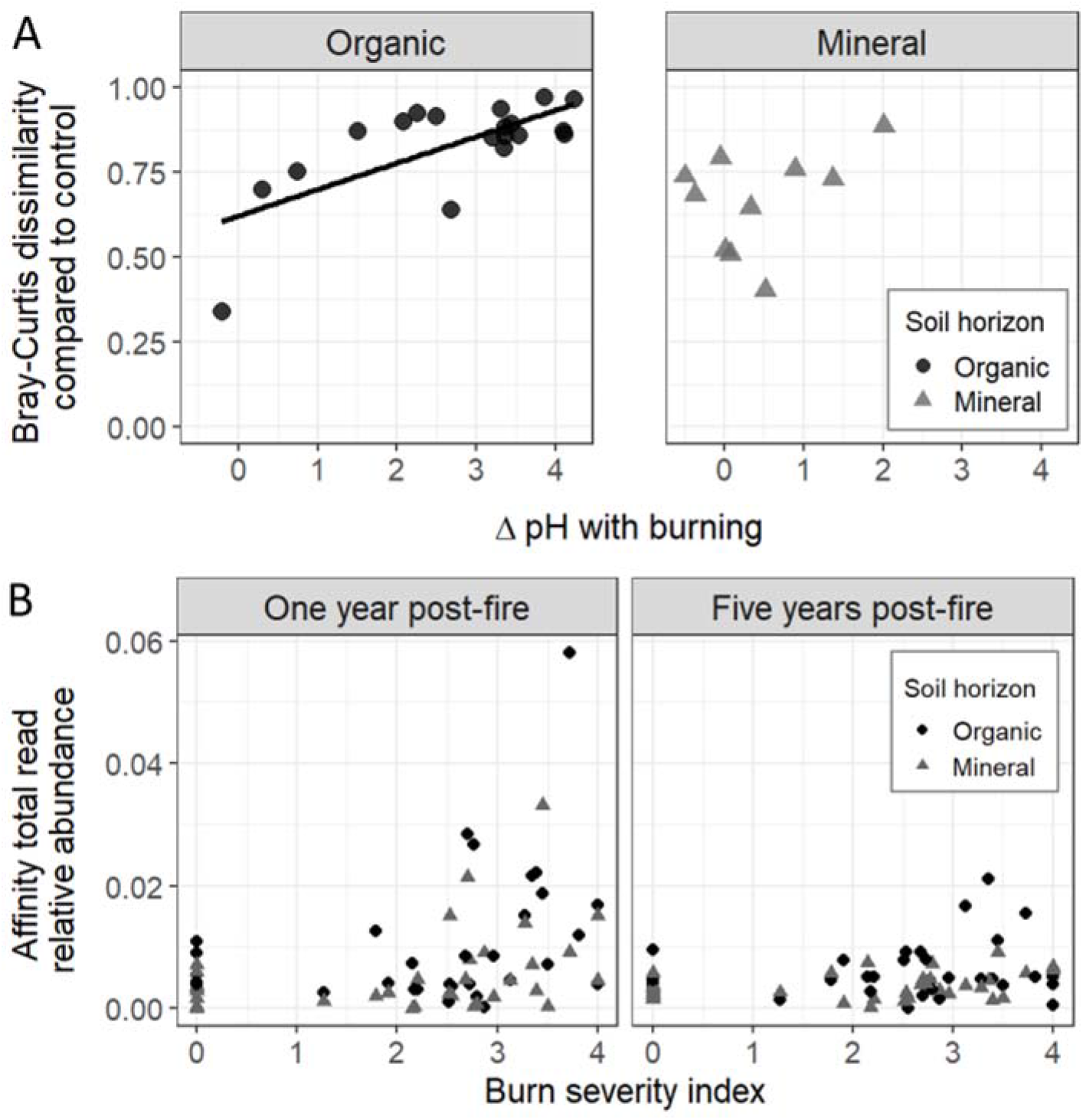
(A) Bray-Curtis dissimilarities in dry soil burn cores compared to unburned controls following autoclaving, inoculation with unburned soil, and a 6-month incubation in organic (left) and mineral (right) horizons vs. change in pH with burning. Line indicates linear model fit (y=0.08x + 0.6; R^2^_adj_=0.50; p=0.0008). (B) Total relative abundance of lab-identified post-fire environment affinity taxa in field data one (left) and five (right) years post-fire, *vs*. burn severity index. Black circles indicate organic horizon samples while grey triangles indicate mineral horizon samples.

38 bacterial OTUs were identified as significantly enriched in burned soil cores compared to unburned control cores, following inoculation with unburned soil and a 6 month incubation (designated “post-fire environment affinity taxa”; Supplementary, Table S11). Of these OTUs, one was also identified as fire surviving and nine were identified as fast growing. These taxa with affinity for the post-fire environment made up a maximum of 6% of the reads in burned communities from the field and were significantly positively correlated with burn severity index (p=0.001, R^2^_adj_=0.11). This relationship was weaker five years post-fire than one year post-fire (interaction term between years post-fire and burn severity index p=0.06).

### Strategic tradeoffs

Only twenty taxa were assigned to more than one strategy, and only one taxon was identified as being significant for all three strategies. For taxa that were significant for at least one strategy (Fig. 7), there were no correlations or only very weak (fast growth and survival, R^2^_adj_=0.03) correlations between strategy strengths, as measured by the log_2_-fold change for each strategy.

**Fig. 7.**
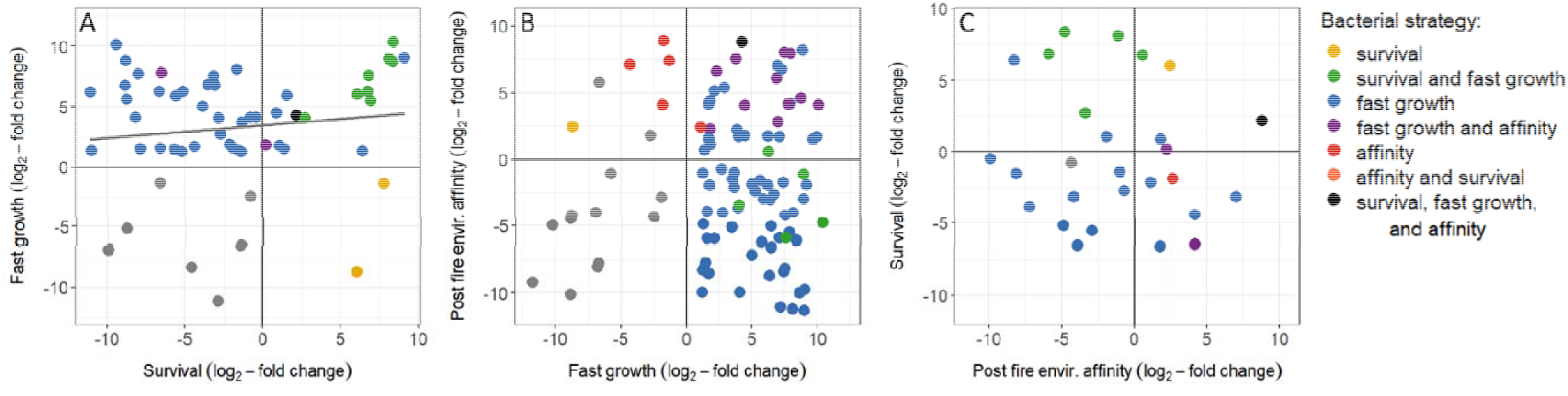
Relative strength of each strategy, as measured by log_2_-fold change in laboratory experiments for (A) fast-growing versus fire-surviving taxa, (B) post-fire environment affinity versus fast-growing taxa, and (C) fire-surviving versus post-fire environment affinity taxa. There was a weak positive relationship between fire survival and fast growth (y=0.2x+1.9; p = 0.03; R^2^_adj_=0.03). Taxa are colored by their assigned strategies.

## Discussion

### Changes in community composition paired with respiration data offer evidence for fire’s effect on C cycling

Our experimental burns directly killed microbes (Fig. 4A) and also altered soil physical and chemical properties (Fig. 2), which, together, caused significant changes to bacterial communities and were characteristic of natural wildfires (Fig. 3). The burn-induced changes to the chemical environment were typical of wildfires - at high temperatures, the complete combustion of organic C and formation of mineral ash has been observed to increase soil pH by as much as 3 units [15]. The decreases in C:N we observed in organic horizons are likely explained by the lower temperature thresholds for C volatilization than for N. During combustion of OM, C losses begin around 100 °C, whereas N volatilization begins around 200 °C [77, 78]. Although narrower C:N ratios are often thought of as a proxy for higher lability, during combustion and PyOM production [32], accompanying chemical changes to soil OM may override this trend [79].

While total C was lost during combustion (Supplementary, Fig. S2), resulting in lower respiration rates on a total dry mass basis (Supplementary, Fig. S7), fire-induced changes in soil properties, combined with direct and indirect effects of fire on microbial communities, also affected mineralization of the remaining C (Fig. 2C; Supplementary, Fig. S8). The decrease in fractional size of the fast C pools with burning (Fig. 2C) is consistent with loss of easily mineralized OM via combustion. The increase in the decay rate of the fast C pool with burning despite the decrease in its fractional size (Fig. 2C) could be explained by a change in the chemical composition such as the production of necromass and increased oxidation of remaining OM [78]. The increase in fractional size and decrease in decay rate of the slow C pools (Fig. 2C) with burning is consistent with pyrolysis of OM during burning leading to production of PyOM, which is less readily decomposed by microbes [35, 80, 81].

We expect that the observed changes in C mineralization were likely driven primarily by changes in the soil environment and changes to the OM chemistry and stocks, rather than by a loss of function in the microbial community. The evidence supporting this interpretation is that inoculating the soil with living microbes did not affect mineralization rates: the C fluxes in the uninoculated 5-week fast growth incubation were indistinguishable from the C fluxes for the first five weeks of the inoculated (post-fire environment affinity) incubations (Supplementary, Fig. S9). If the microbial community had been significantly impaired with respect to its ability to mineralize C, due to either changes in composition or due to persistent reductions in biomass, we would have expected inoculation to increase C mineralization rates, which we did not observe. Microbes can rapidly grow in the post-fire environment, as evidenced by the rapid proliferation of fast growers (Fig. 5).

### Importance of fire survival is small but can persist for years

We hypothesized that fire survival would be most important in structuring bacterial communities immediately post-fire and following higher-temperature burns, which was generally supported by our data (Fig. 4). Unsurprisingly, burning caused a decrease in total RNA and DNA concentrations in the soil (Fig. 4A), which can be explained by bacterial (and other organismal) mortality, and destruction of necromass and relic DNA [82, 83]. However, our approach to identifying fire survivors yielded only a very small number of taxa as fire survivors (17 of 15079 total OTUs across the laboratory dataset), suggesting, somewhat counter-intuitively, that fire survival is not an important strategy, even immediately following a fire.

How can that be? We posit that there is likely a relatively narrow optimal zone [84] for fire temperatures where survival plays a meaningful role. At sufficiently high burn temperatures, there is likely almost complete microbial death [17]. However, at low burn temperatures, characteristic of wet soil burns and deeper soil horizons (Fig. 2A), there is very little microbial death (Fig. 4A), so survival is again not a relevant strategy. This observation is supported by the field data, where the sites the greatest proportion of fire survivors had intermediate severity burns (Fig. 4B).

In the field, any short-term effects of fire survival may also be rapidly overwhelmed by dispersal, which can increase soil bacterial community resilience [85, 86]. In boreal forests, spatial heterogeneity in fire severity results in complete combustion of soil O horizons in some areas while adjacent “fire refugia” remain unburned [87, 88]. Similarly, even when lethal burn temperatures are reached at the surface, temperatures rapidly attenuate with depth (Fig. 2A), supporting belowground fire refugia from which dispersal to surface horizons could occur as well. Thus, we propose that fire survival can be an important strategy structuring post-fire microbial communities after certain fires and timescales, but only when a narrow range of conditions is met: (1) temperatures reach levels that are lethal for some taxa *but not for others*; (2) post-fire dispersal from above or below is not sufficient to overwhelm this effect.

For the few taxa identified as fire-survivors, what traits allow them to survive? Fire survival may be associated with dormancy [20, 89]. Some *Actinobacteria* and *Firmicutes* enriched following high severity burns have been found to possess sporulation genes, suggesting that sporulation and dormancy may aid in fire survival [90, 91]. One of our fire survivors was classified as *Bacillus sp*., which is a known endospore-forming genus [92], and accounted for up to 8% of total reads in dry soil burns. Taxa such as these may survive burning in a dormant state and then either germinate and grow, or persist in a dormant state after the fire. Some taxa enriched following high severity burns have also been found to possess genes encoding heat shock proteins and/or a higher GC content, both of which may be additional traits promoting fire survival [90]. About half of fire survivors (9 of 17), including *Bacillus* sp., were also identified as being fast growers, highlighting potential positive interactions between strategies and the possibility of dormant cells surviving and germinating post-fire, and then rapidly colonizing the burned soil. However, fire survival was a relatively uncommon strategy compared to fast growth, and the traits promoting fire survival – such as dormancy and heat shock proteins – are not necessarily expected to directly promote fast growth.

### Importance of fast growth is large and short-lived

We hypothesized that, with time, the importance of fire survival would be superseded by fast growth, as fast growth would enable bacteria to take advantage of an increase of available nutrients released via burning and rapidly colonize the burned environment. Our results were generally consistent with this hypothesis, as the number of taxa identified as having potential for fast growth (3% of OTUs) far exceeded that of fire-surviving taxa (0.1% of OTUs). Although burned soils were not initially enriched in putative fast-growers, by five weeks post-burn, they had higher weighted predicted mean 16S rRNA gene copy numbers than unburned controls (Fig. 5A), indicating that fast growth quickly becomes important following burning. This rapid increase in fast-growing taxa is likely fueled by nutrients liberated by fire via heat-induced bacterial mortality, vegetation death, and pH increase, combined with reduced competition [24]. In the field, these fast-growing taxa account for a large (up to 40%) of total reads one year post-fire, and were positively correlated with burn severity one-year post-fire (Fig. 5B), further indicating that – in the short-term – fast growth plays a strong role in structuring community composition. However, the positive correlation between burn severity and relative abundance of fast growers completely disappears by five years post-fire. Despite this, microbial communities can take longer than five years to recover post-fire (Fig. 3; 94,95), indicating that strategies other than fast growth likely play key roles in structuring soil communities over decadal timescales.

Are there important implications for ecosystem functioning due to this bloom of fast-growing taxa? For the first two weeks, total respiration rates (per gram initial soil C) were equivalent or higher in dry burned soil than in unburned soil (Supplementary, Fig. S8). This suggests that any limitation to mineralization due to high initial bacterial mortality (Fig. 3A) was rapidly offset by fast growing taxa. However, by three weeks post-burn, total respiration rates (per gram initial soil C) were lower in dry burned soil than in unburned soil (Supplementary, Fig. S8). This reversal in trend could indicate that microbes rapidly depleted the already smaller fast C pool, shifting to greater reliance on the larger but more persistent fire-altered slow C pool (Fig. 2C).

### Importance of post-fire environment affinity is small and relatively short-lived

We hypothesized that in the months to years following fire, an affinity for the post-fire soil environment (*i.e.*, an affinity for post-fire soil conditions and fire-generated compounds) would be the most important strategy structuring bacterial communities. While this was generally supported in our incubation study, the importance of this strategy over longer timelines in the field was small (Fig. 6B). This may be due in part to the fact that “affinity for the post-fire soil environment” likely encompasses myriad traits that depend on specific soil properties and their response to a given burn. That said, there may be some properties that tend to be common across burned soils.

Physicochemical changes to soil properties such as pH post-fire were often extreme (Fig. 2B) and likely constrained post-fire community composition, offering opportunities for bacteria with adaptive traits to thrive. The soils where the community changed the most with burning also had the greatest increases in pH (Fig. 6A), which could help explain why reintroduction of bacteria from unburned soils had negligible effects on community composition.

Along with pH shifts, post-fire environment affinity could be conferred by the ability to decompose PyOM produced during the fire. While we didn’t measure PyOM directly, all dry soil burns resulted in some degree of organic horizon combustion, and temperatures during the dry soil burns would have been high enough to produce PyOM [95]. This is supported by the increase in the fractional size and decrease in decay rate of the slow C pool of the burned soils, as soil C in the form of PyOM would be expected to be represented within the slow C pool (Fig. 2C). The ability to degrade PyOM may become increasingly important after easily-mineralizable fire-liberated C, such as dead microbes and fine roots, is depleted but before fresh inputs from vegetation have returned to pre-fire levels. Supporting this, in the field data, the importance of post-fire environment affinity taxa was positively correlated with burn severity (Fig. 6B), and this relationship weakened five years post-fire, possibly driven by fresh inputs to fast C pools returning as plant communities rebounded.

Post-fire environment affinity taxa make up a smaller proportion of the total community in the field (up to 6%) than in the lab (up to 20%), where soil communities have been isolated from the potential effects associated with plant recolonization. Thus, for determining microbial community composition, we might propose that the immediate effects of fire on soil properties, such as pH shifts or losses and transformations of soil OM, are likely to be less important than the effects on soil conditions mediated by plant community re-establishment that emerge over time such as changes to C inputs, water and nutrient availability, symbioses, and microclimate effects [96].

### Strategy-based framework

Our strategy-prediction approach identified taxa representing up to 60% of total reads in laboratory-burned samples and 45% in field data (Supplementary, Fig. S10). However, most responsive taxa (97%) were only assigned to one strategy (Fig. 7). On the one hand, this paucity of OTUs associated with two or more strategies could offer evidence for a trade-off in strategies. However, for taxa with multiple strategies, a strategic tradeoff should manifest as a negative relationship between the strength of different strategies, which we did not observe (Fig. 7). This lack of evidence for strategic trade-offs is somewhat counter to our and others’ previous conceptions of post-fire survival strategies as fitting into a framework analogous to the Yield-Acquisition-Stress-tolerant framework [8, 97]. While we would not argue that the results in this paper offer conclusive evidence that trade-offs are not relevant for post-fire success strategies, we note that some fire-relevant strategies may be complementary. For example, *Bacillus* sp. are often noted both for their ability to form stress-resistant endospores and also for their high 16S rRNA gene copy number and potential for fast growth [26, 98]. Further work will be needed to fully explore the prevalence of strategy co-occurrences or trade-offs post-disturbance.

Drawing on our observations, we offer a hypothetical framework of response to fire for bacterial communities (Fig. 8). The importance of the three fire-adaptive strategies presented here varies over time and with burn severity (or soil depth).

**Fig. 8.**
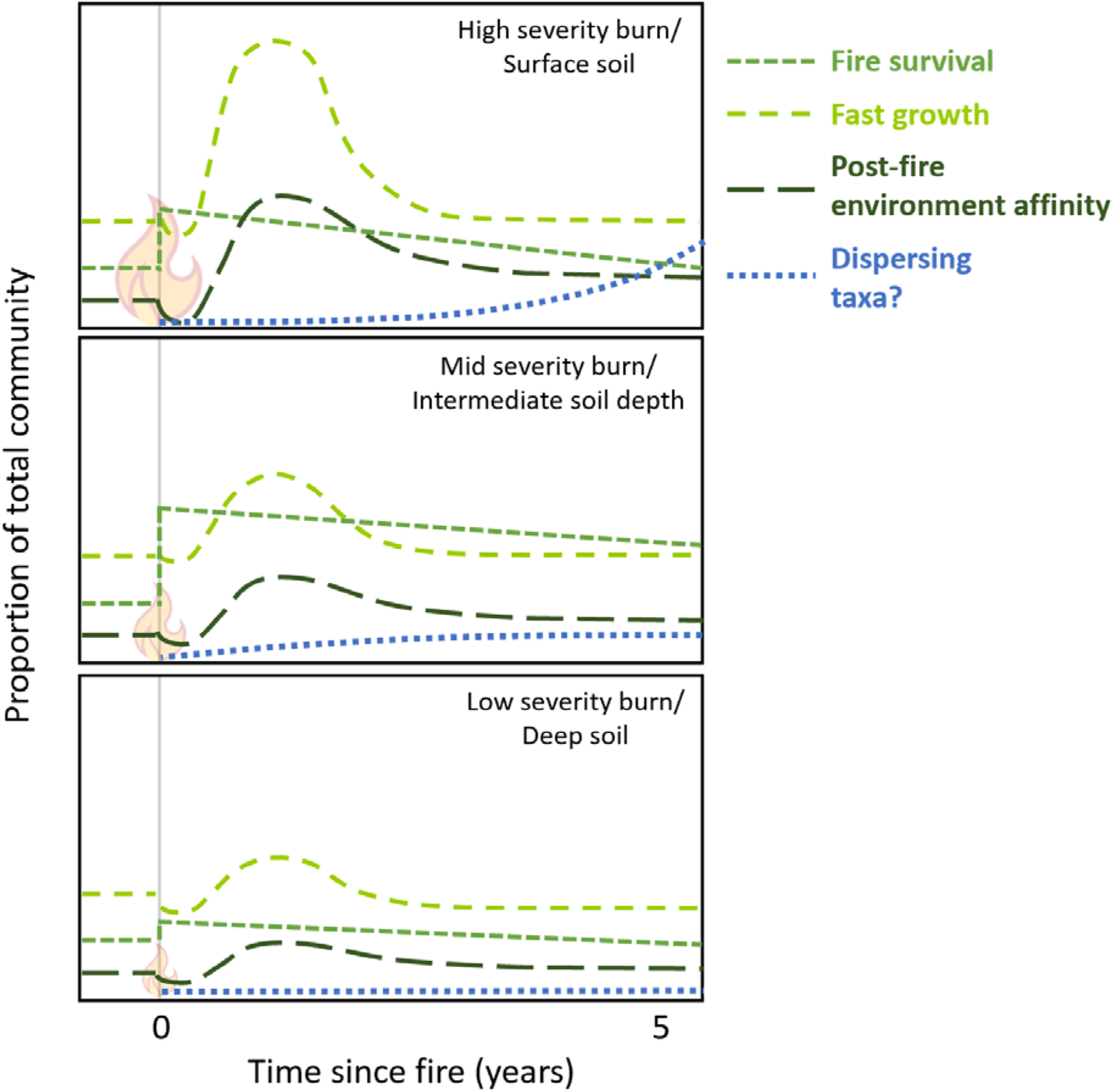
Hypothetical framework showing the relative importance of fire survival, fast growth, post-fire environment affinity, and dispersal in structuring post-fire soil bacterial communities across high (top panel), mid (middle panel), and low (bottom panel) severity burns or soil depths. (Note that, while hypothesized fire survival, fast growth, and post-fire environment affinity trends are based directly on our observations in this study, our predictions for the importance of dispersing taxa are largely speculative.)

Immediately post-fire, some taxa are particularly strong survivors. However, this effect is primarily relevant for burns of intermediate severity and/or intermediate (∼50°C-100°C) soil temperatures, with lower-temperature burns leaving the community generally intact, and higher-temperature burns wiping out all organisms. Under these survivor-optimal burn conditions, fire-surviving taxa can remain enriched in the community for years after the fire (Fig. 4B). Fast growing bacteria rapidly fill the niche created by bacterial mortality and a pulse of available C sources, making up large portions of the total community. Higher severity burns select for more fast-growing taxa due to greater overall mortality and a larger open niche space. However, the rapid proliferation of these taxa essentially disappears between one and five years post-fire. Over the same time period, a smaller set of taxa with an affinity for post-fire environmental conditions thrive, possibly via tolerating fire-induced changes in soil properties such as pH, or by specialization for fire-generated compounds. Due to above- and belowground fire refugia, dispersal limitation may not be a particularly important factor in structuring post-fire communities shortly after the fire, even for high-severity burns, and establishment of dispersing organisms may be constrained by fire-affected chemical properties. Over longer periods, additional strategies not characterized in this study, such as those related to the effects of recolonizing plant community on soil properties [96], may become increasingly relevant.

### Conclusion

Using simulated burns and laboratory incubations to identify fire-surviving taxa, fast-growing taxa, and post-fire environment affinity taxa, we were able to assign strategies for up to 45% of total reads in field samples collected across a range of vegetation and soil types and offer a hypothetical framework for the relevance of these post-fire strategies. Further work remains to explore the functional implications of these strategies, identify additional strategies not characterized in this study, and evaluate relationships between recolonizing plant communities, soil properties, and microbial strategies over decadal timelines. Next steps will also include testing this framework and identifying these taxa in other fire-adapted ecosystems.

## CRediT Author Statement

**Dana Johnson:** Conceptualization, Methodology, Software, Formal analysis, Investigation, Writing – Original draft, Writing – Review and editing, Visualization. **Jamie Woolet:** Methodology, Investigation, Writing – Review and editing. **Kara Yedinak:** Methodology, Investigation, Resources and Data curation, Writing – Review and editing. **Thea Whitman:** Conceptualization, Methodology, Software, Formal analysis, Investigation, Writing – Review and editing, Supervision, Project administration, Funding acquisition.

## Acknowledgements

This work was funded by the University of Wisconsin-Madison Fall Research Competition Grant and the U.S. Department of Energy (DE-SC0021022). We thank Dr. Ellen Whitman for her assistance in identifying field sites and burn history and for support in field collection. We thank Dr. Daniel K. Thompson for his insights on burn characteristics for this region. We thank Laura Hasburgh and Keith Bourne at the USDA FS Forest Products Laboratory for their advice and assistance with burn simulations. We thank Jean Morin, Sharon Irwin, and other Wood Buffalo National Park staff for their support in conducting this research (Permit WB-2019-31497).

## Competing Interests

The authors declare no competing interests.

## Data Availability Statement

The sequencing datasets generated during the current study are available in the NCBI SRA under bioproject number [SUB11346170]. Field sequencing data are available at PRJNA564811 (2015 data) and PRJNA825513 (2019 data, available 31 December 2022). Non-sequencing data are deposited in the DOE ESS-DIVE repository under: https://data.ess-dive.lbl.gov/datasets/ess-dive-e00cfbe3d89f2c1-20220421T224630867

Code for the analyses conducted in the current study are available at https://github.com/DanaBJohnson/WoodBuffalo.

## Notes

https://data.ess-dive.lbl.gov/datasets/ess-dive-e00cfbe3d89f2c1-20220421T224630867

https://github.com/DanaBJohnson/WoodBuffalo

https://www.ncbi.nlm.nih.gov/bioproject/PRJNA564811/

https://www.ncbi.nlm.nih.gov/bioproject/PRJNA825513/

### Competing Interest Statement

The authors have declared no competing interest.

