## Supplementary Information for "An empirical approach to developing and testing a traits-based fire ecology framework for bacterial response to wildfires"

Table of Contents	Page

**Supplemental Figures**

1. Soil collection and burning photos	3
2. Total C and N	3
3. Two-pool decay model fits for the fast-growth incubation	4
4. Two-pool decay model fits for the post-burn affinity incubation	5
5. Post-burn affinity incubation C pool fractional size and turnover rate	6
6. NMDS ordination on weighted UniFrac of genomic DNA across all experiments	7
7. Respiration rate per initial gram dry soil for the post-burn affinity incubation	8
8. Respiration rate per initial gram C for the post-burn affinity incubation	9
9. Comparison of respiration rate between inoculated and uninoculated incubations	10
10. The percent of total community identified as taxa with a fire-adaptive strategy	11

**Supplemental Tables**

1. Site characteristics, location, and soil properties	12
2. Mean fractional dry mass loss	13
3. Mean soil temperatures during simulated burns	14
4. Mean soil pH, total C, total N, and horizon thickness	15
5. Mean 2-pool decay coefficients for fast-cycling C pool post-burn	16
6. Mean 2-pool decay coefficients for slow-cycling C pool post-burn	16
7. PERMANOVA results for full model for 16S for cDNA	17
8. PERMANOVA results for full model for 16S for genomic DNA	17
9. PERMANOVA results for full model at end of fast-growth incubation	18
10. PERMANOVA results for full model at end of post-fire affinity incubation	19
11. List of all OTUs identified for each fire-adaptive strategy	19

**Supplemental Notes	20**

**References	23**

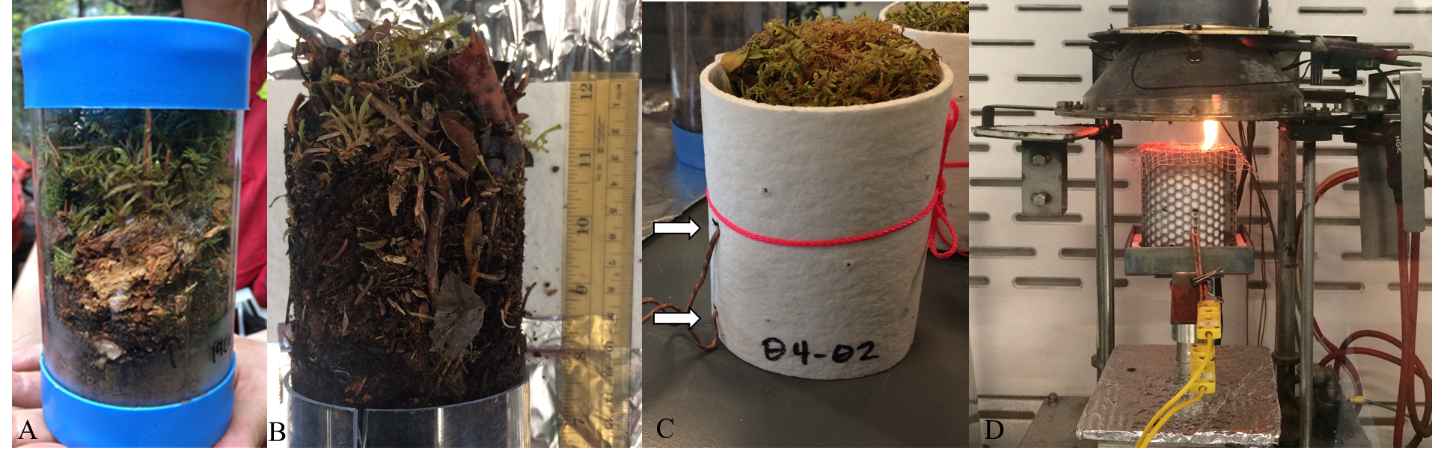

Figure S1. Soil cores (A) collected, (B) extruded, (C) wrapped in ceramic paper in preparation for the burn. Arrows indicate the position of the mid and lower thermocouples within the core. (D) Soil cores exposed to heat flux in cone calorimeter.

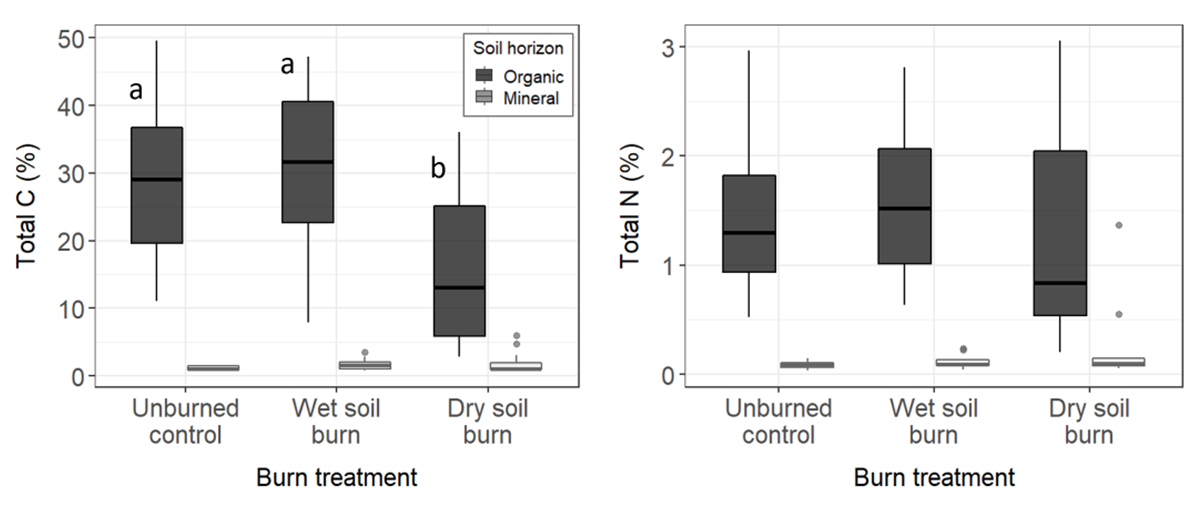

Figure S2. Soil total C (left) and total N (right) across the three burn treatments. Dark fill indicates organic horizon samples while light grey indicates mineral horizons. Different letters indicate significant differences between treatments for a given horizon.

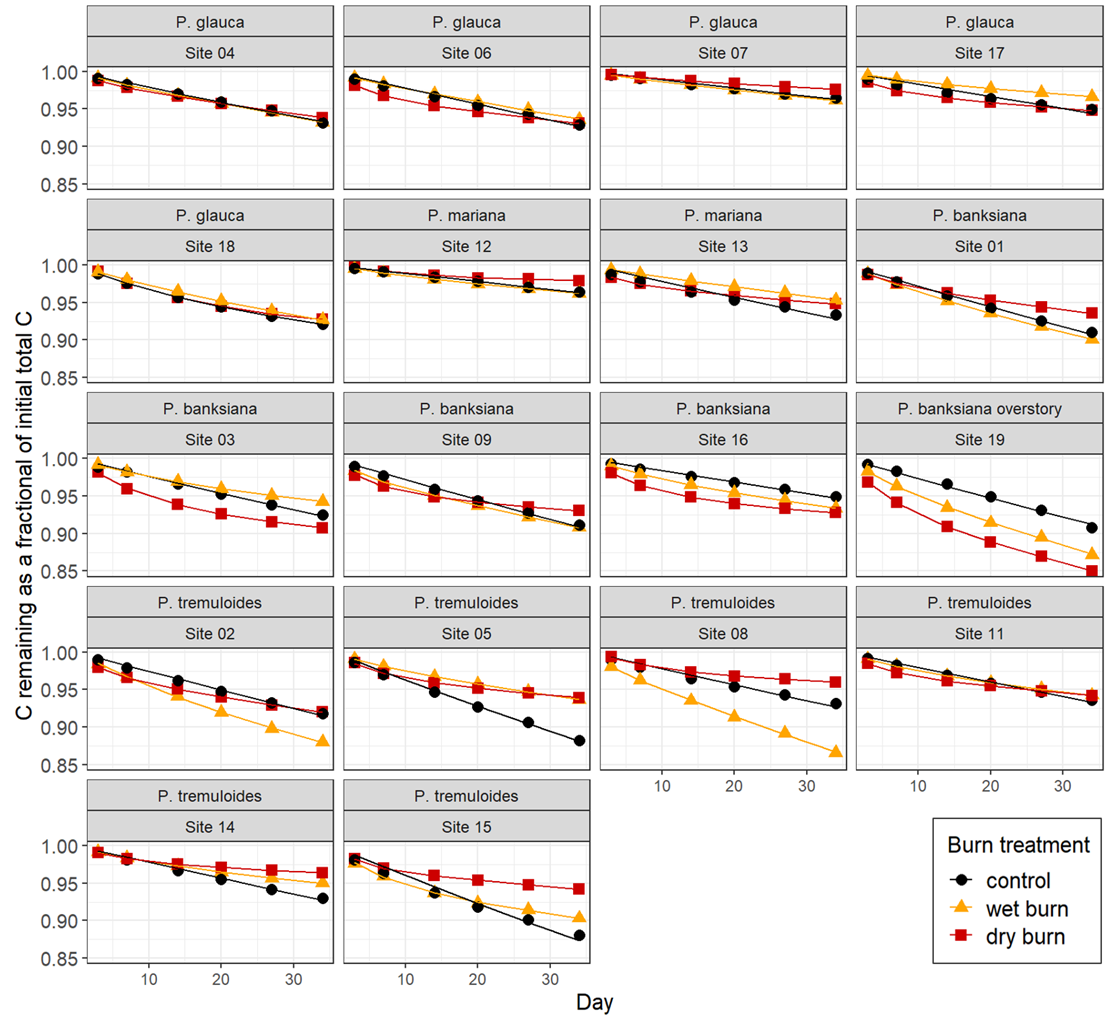

Figure S3. Carbon remaining as a fraction of initial total C in burned and unburned soil across all sites with 2-pool decay model fits for the 5-week fast-growth incubation.

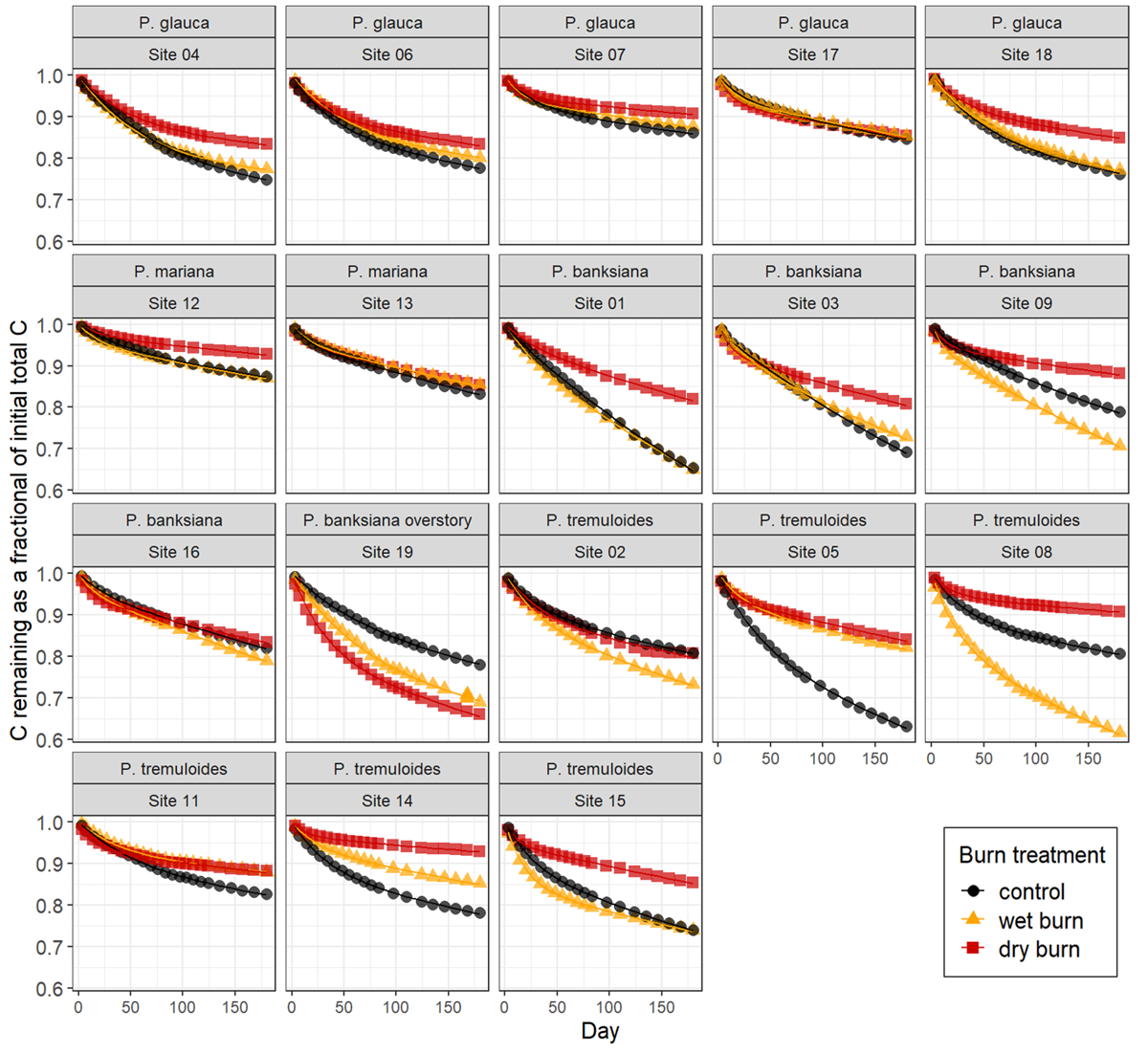

Figure S4. Carbon remaining as a fraction of initial total C in burned and unburned soil across all sites with 2-pool decay model fits for the 6-month post-fire affinity incubation following autoclaving and inoculation with unburned soil.

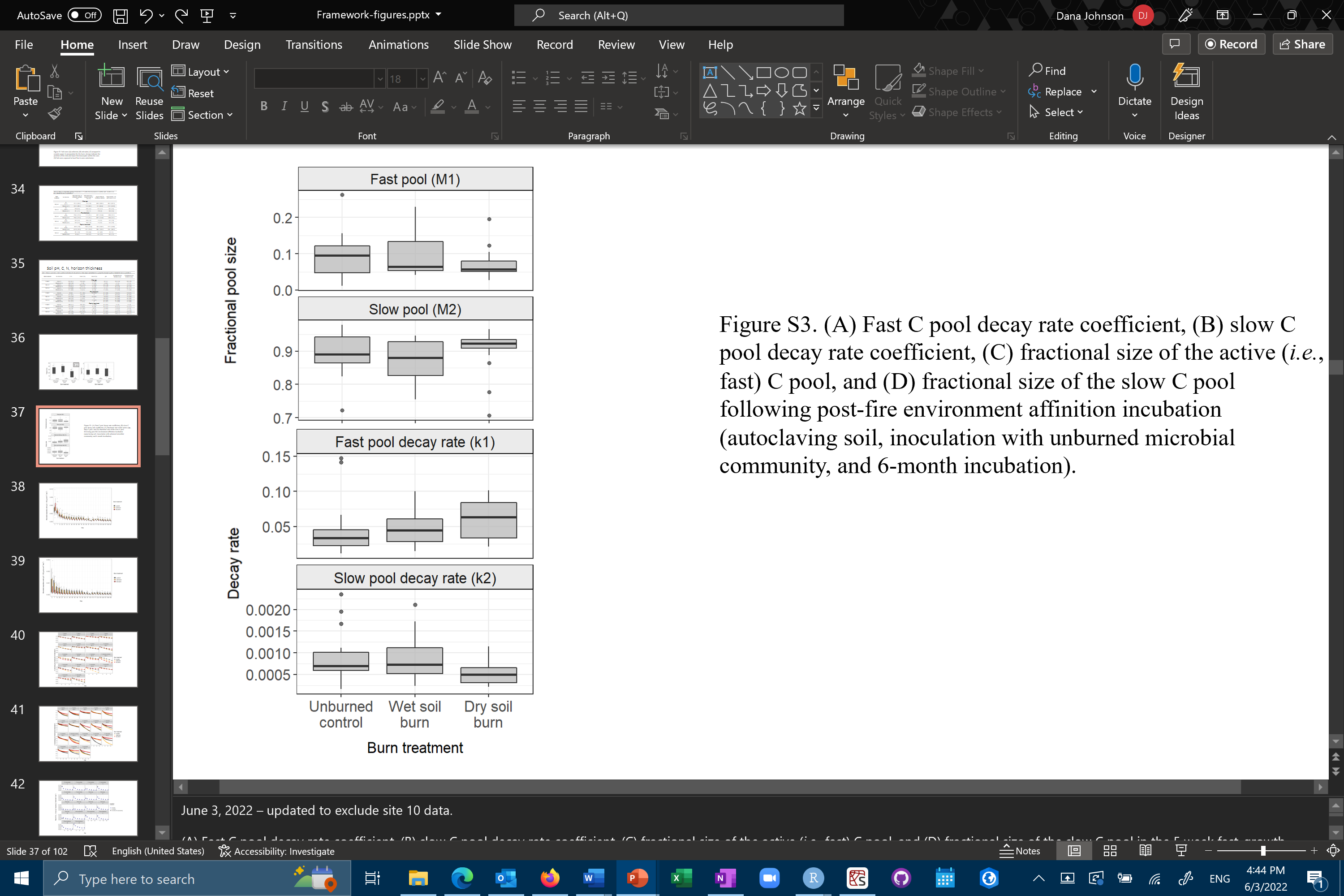

Figure S5. (A) Fractional size of the fast (*M_1_*) and slow (*M_2_*) C pool (top two panels) and decay rate coefficients for the fast (*k1*) and slow (*k2*) C pool (bottom two panels) of the slow C pool following post-fire environment affinition incubation (autoclaving soil, inoculation with unburned microbial community, and 6-month incubation).

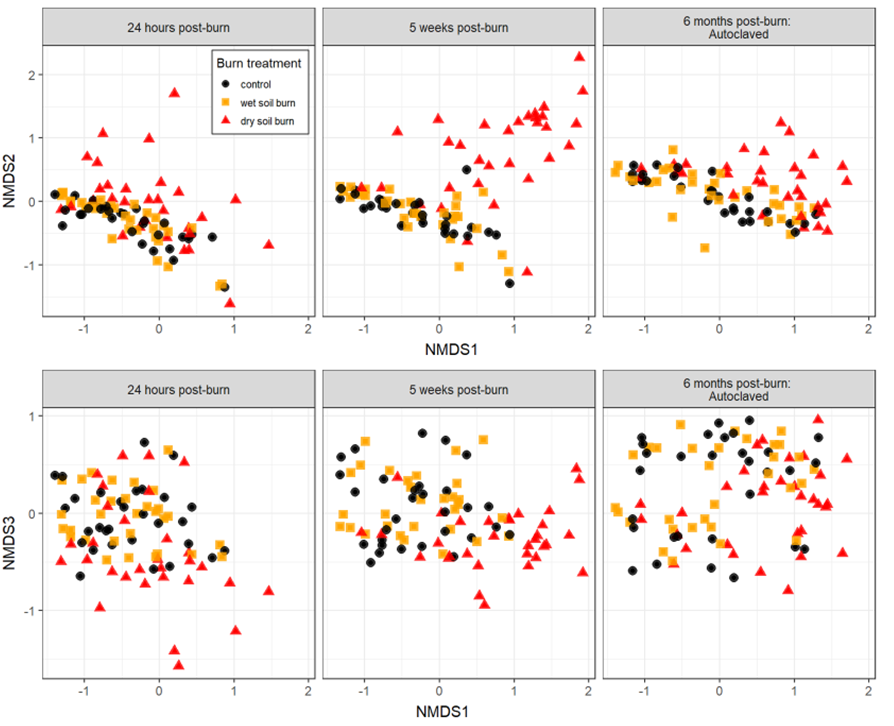

Figure S6. NMDS ordination on weighted UniFrac of genomic DNA rRNA genes across all experiments (k=3, stress=0.12). One ordination was performed and is faceted by the three experimental timepoints– 24 hours, 5 weeks, and 6 months (following autoclaving and inoculation with unburned soil). Black, gold, and red indicate unburned control, wet soil burned, and dry soil burned, respectively. Circles, squares, and triangles indicated unburned, wet soil burned, and dry soil burned, respectively.

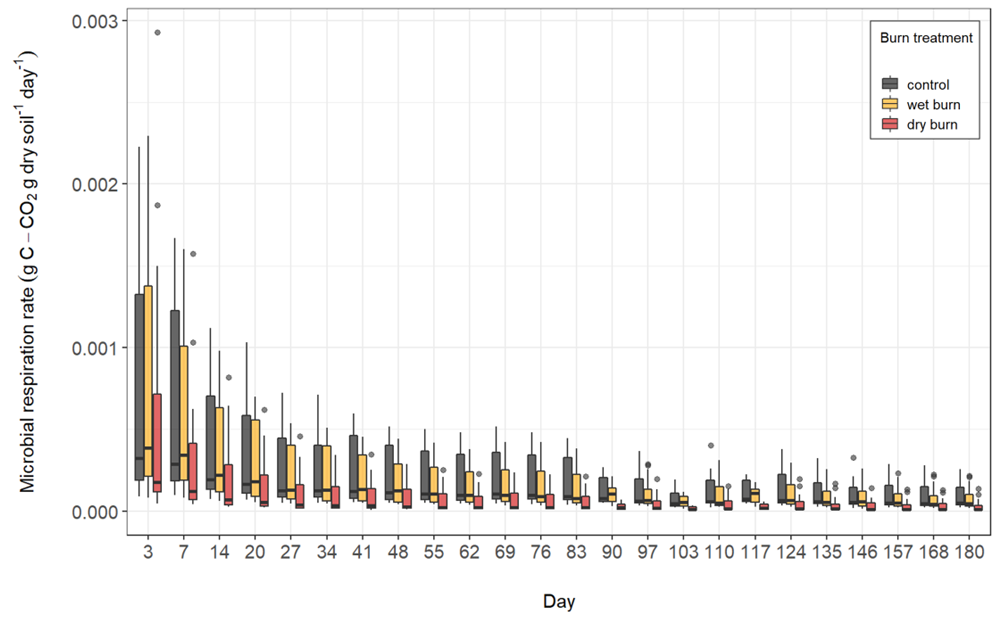

Figure S7. Respiration rate per initial gram dry soil for unburned (grey), wet soil burn (yellow), and dry soil burn (red) from each site over the course of the 6-month incubation following autoclaving and inoculation with unburned soil.

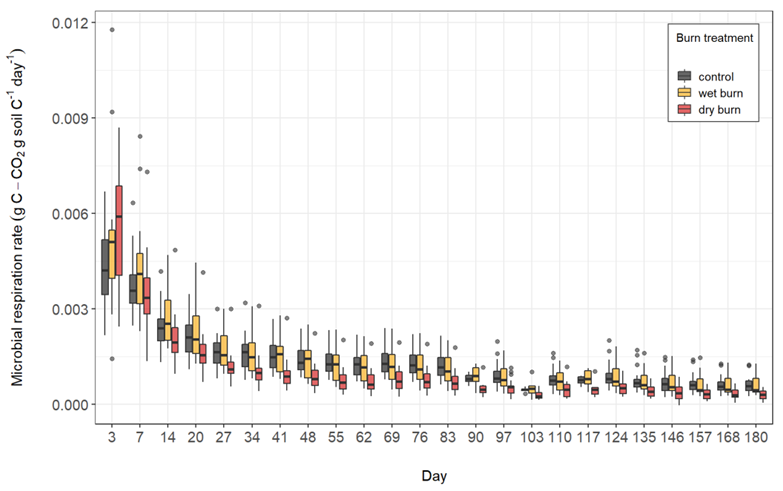

Figure S8. Respiration rate per initial gram C for unburned (grey), wet soil burn (yellow), and dry soil burn (red) from each site over the course of the 6-month incubation following autoclaving and inoculation with unburned soil.

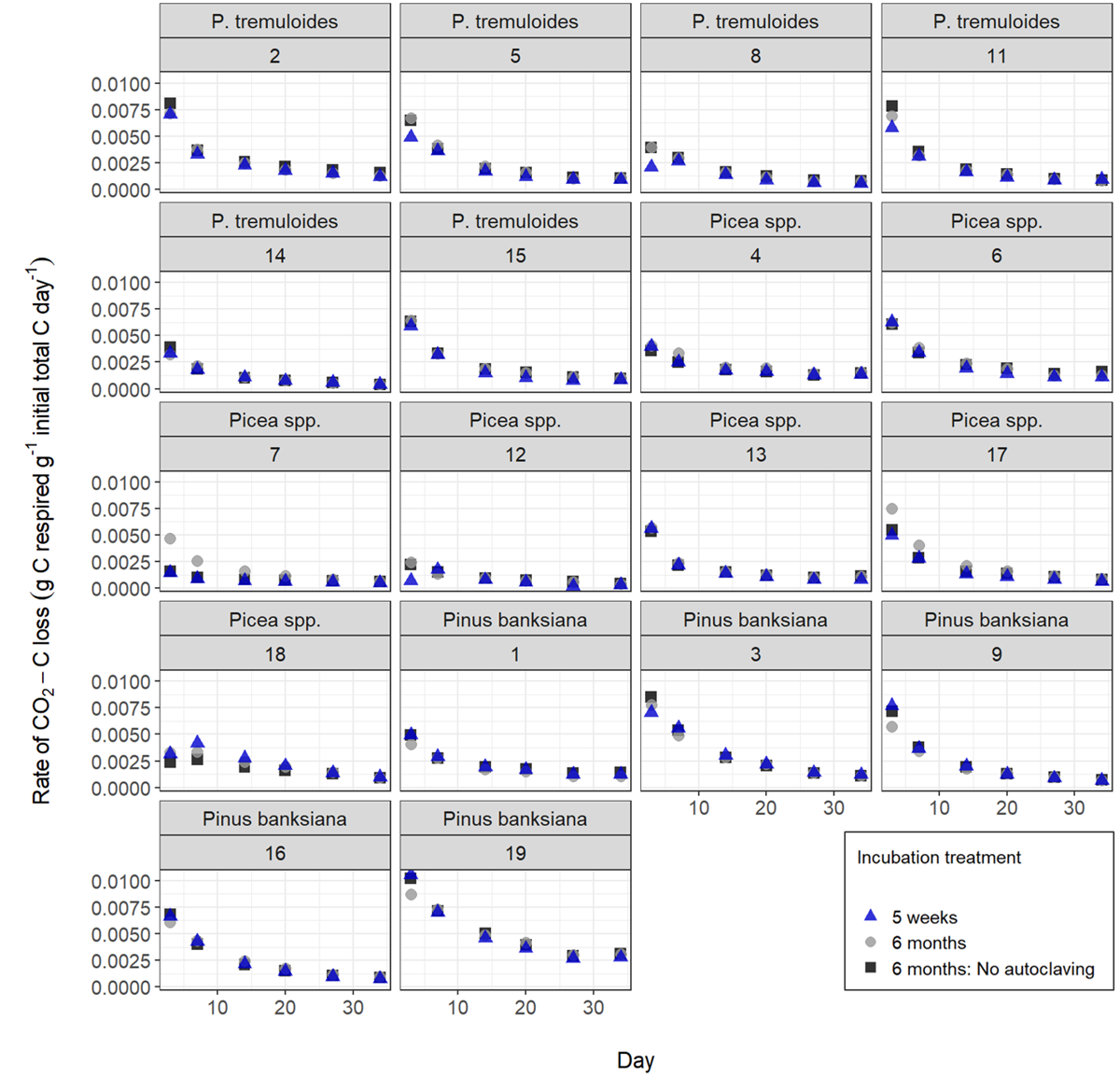

Figure S9. Comparison of respiration rate (per initial gram C) for dry burn soil over the first 5 weeks of the 5-week fast growth incubation (blue), 6-month post-fire environment affinity incubation including autoclaving and inoculation with unburned soil microbial community (grey), and 6-month incubation including inoculation but excluding autoclaving (black).

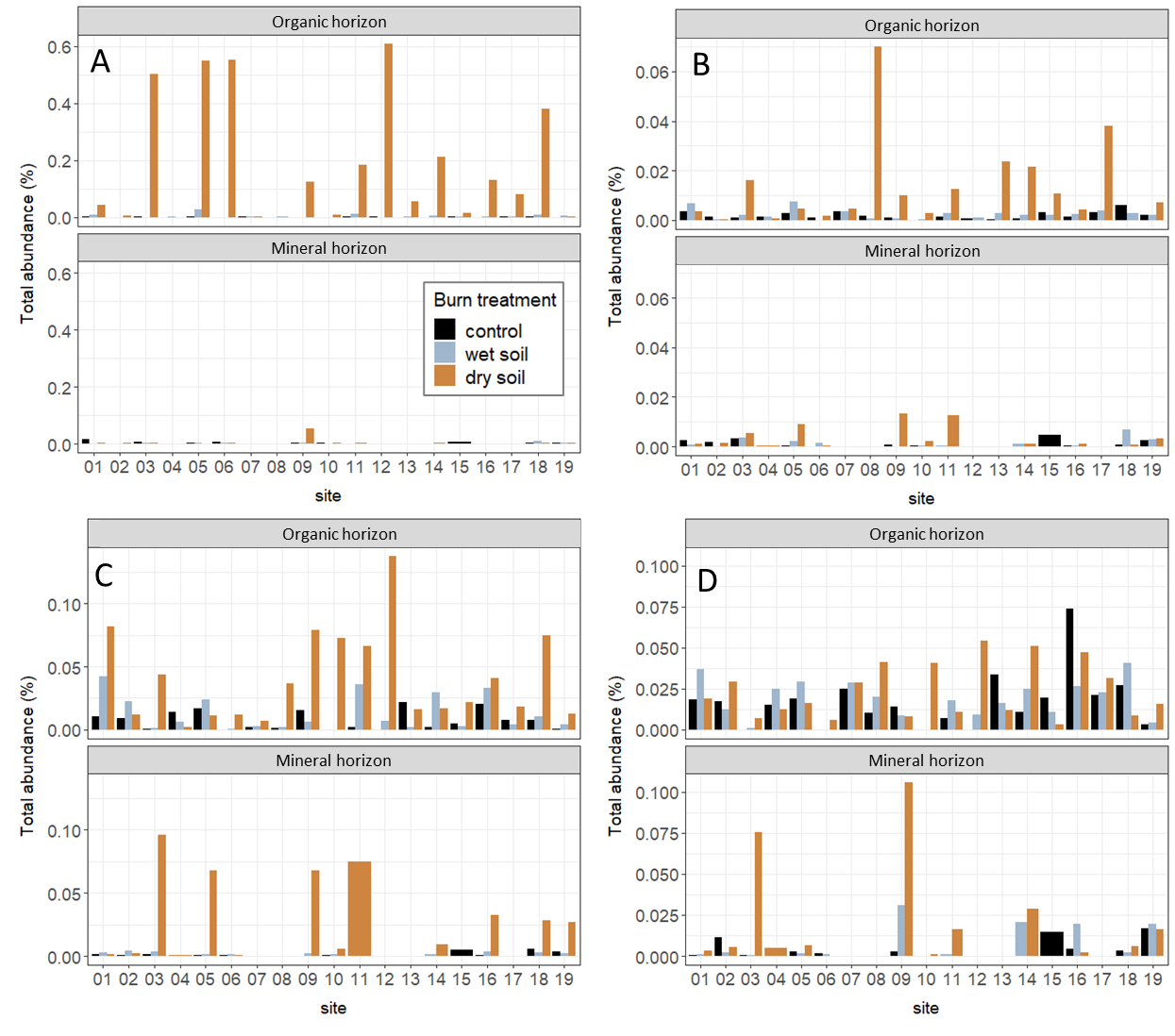

Figure S10. The percent of total reads identified as taxa with any of the three fire-adaptive strategies (A) RNA and (B) DNA samples immediately post-fire, (C) 5-week fast growth incubation, and (D) 6-month post-fire environmental affinity incubation following autoclaving and inoculation with unburned soil.

| Table S1. Site characteristics, location, and soil properties grouped by dominant vegetation type. | | | | | | | | |
| --- | --- | --- | --- | --- | --- | --- | --- | --- |
| Site | Slope position | Aspect | Overstory density (%) | Latitude | Longitude | Sand (%) | Silt (%) | Clay (%) |
| ***Picea* spp.** | | | | | | | | |
| 4 | Middle | ESE | 98 | 59° 27' 44.62" N | 112° 17' 51.10" W | 62.5 | 32.5 | 5 |
| 6 | Lower | NW | 52 | 59° 28' 19.14" N | 112° 16' 23.58" W | 70.5 | 19.5 | 10 |
| 7 | Level | NNW | 96 | 59° 35' 29.05" N | 112° 16' 13.72" W |  | NA |  |
| 12 | Toe |  | 91 | 59° 48' 28.97" N | 112° 00' 18.41" W |  | NA |  |
| 13 | Level |  | 79 | 59° 58' 24.99" N | 112° 26' 57.74" W |  | NA |  |
| 17 | Level |  | 92 | 60° 01' 48.23" N | 112° 53' 21.29" W |  | NA |  |
| 18 | Level |  | 92 | 60° 00' 54.13" N | 112° 52' 42.88" W | 33 | 37 | 30 |
| ***Populus tremuloides*** | | | | | | | | |
| 2 | Level |  | 94 | 59° 27' 07.18" N | 112° 19' 35.98" W | 54.5 | 32 | 13.5 |
| 5 | Middle | NW | 97 | 59° 27' 41.18" N | 112° 17' 00.31" W | 59.5 | 31 | 9.5 |
| 8 | Level | S | 99 | 59° 38' 59.28" N | 112° 12' 45.89" W | 68 | 25 | 7 |
| 11 | Level |  | 99 | 59° 47' 45.65" N | 112° 02' 27.45" W | 59 | 32 | 9 |
| 14 | Level |  | 89 | 60° 02' 54.21" N | 112° 47' 50.94" W | 72 | 20 | 8 |
| 15 | Lower | NW | 93 | 60° 02' 21.03" N | 112° 52' 45.09" W | 70 | 19 | 11 |
| ***Pinus banksiana*** | | | | | | | | |
| 1 | Upper | W | 96 | 59° 24' 38.19" N | 112° 23' 50.15" W | 23 | 43 | 34 |
| 3 | Level |  | 100 | 59° 27' 25.16" N | 112° 19' 27.27" W | 51.5 | 36 | 12.5 |
| 9 | Level |  | 97 | 59° 41' 04.03" N | 112° 10' 18.51" W | 54 | 29.5 | 16.5 |
| 10 | Level |  | 100 | 59° 42' 04.70" N | 112° 10' 13.50" W | 49 | 41 | 10 |
| 16 | Level |  | 85 | 60° 01' 17.15" N | 112° 58' 14.23" W | 50 | 32.5 | 17.5 |
| 19 | Middle | SSE | 91 | 60° 02' 05.49" N | 113° 08' 40.44" W | 93 | 3 | 4 |

| Table S2. Mean whole-core fractional dry mass loss during burning grouped by dominant vegetation type and burn treatment. Standard deviation in parentheses. | | |
| --- | --- | --- |
| **Dominant Vegetation** | **Burn treatment** | **Fractional dry mass loss** |
| *Picea* sp. | dry | 0.2 ± (0.19) |
| *Picea* sp. | wet | 0.14 ± (0.07) |
| *Pinus banksiana* | dry | 0.09 ± (0.05) |
| *Pinus banksiana* | wet | 0.06 ± (0.03) |
| *Populus tremuloides* | dry | 0.23 ± (0.21) |
| *Populus tremuloides* | wet | 0.1 ± (0.03) |

| Table S3. Mean soil temperature and degree hours above 21 °C at the O horizon-mineral soil interface and 1 cm above core base. Standard deviation in parentheses. | | | | | |
| --- | --- | --- | --- | --- | --- |
| Burn treatment | Soil horizon | Maximum temp. at O/Mineral interface (°C) | Maximum temp. 1 cm above base of core (°C) | Degree hours at O/Mineral interface | Degree hours 1 cm above base of core |
| ***Picea* spp.** | | | | | |
| Dry soil | O (n=7) | 357.3 (234.1) | 152 (179) | 699.3 (545.3) | 242.2 (297.5) |
|  | Mineral (n=3) | 292.3 (249.1) | 75.9 (68.2) | 521 (606.4) | 135.2 (163.8) |
| Wet soil | O (n=7) | 29.9 (6.3) | 28.6 (3.3) | 19.3 (7.2) | 26.3 (7.6) |
|  | Mineral (n=2) | 28.1 (2.7) | 29 (1.7) | 25.8 (0) | 34.9 (5.4) |
| ***Pinus banksiana*** | | | | | |
| Dry soil | O (n=6) | 359.6 (157.6) | 71.5 (35.1) | 487.5 (314.4) | 108.4 (53.3) |
|  | Mineral (n=6) | 359.6 (157.6) | 71.5 (35.1) | 487.5 (314.4) | 108.4 (53.3) |
| Wet soil | O (n=6) | 30.5 (5.3) | 26.8 (1.9) | 15 (5.5) | 19.2 (7.8) |
|  | Mineral (n=6) | 30.5 (5.3) | 26.8 (1.9) | 15 (5.5) | 19.2 (7.8) |
| ***Populus tremuloides*** | | | | | |
| Dry soil | O (n=6) | 439.5 (159) | 199.6 (210.4) | 745.9 (396.5) | 323.5 (334.8) |
|  | Mineral (n=4) | 413.8 (195.9) | 176.1 (230.5) | 584.1 (309.3) | 224.1 (246.5) |
| Wet soil | O (n=6) | 28.1 (3.4) | 25.9 (0.4) | 16.7 (4.8) | 20.2 (7) |
|  | Mineral (n=4) | 28 (4.1) | 25.8 (0.5) | 14.3 (3.9) | 18.5 (7.5) |

| Table S4. Mean soil pH, total C, total N, and horizon thickness for dry and wet soil burn samples and unburned soil grouped by dominant vegetation. Standard deviation in parentheses. | | | | | | | |
| --- | --- | --- | --- | --- | --- | --- | --- |
| Burn treatment | Soil horizon | C:N | Total C (%) | Total N (%) | pH | Pre-burn horizon thickness (cm) | Post-burn horizon thickness (cm) |
|  | ***Picea* spp.** | | | | | | |
| Control | O  (n=7) | 22.6 (6.1) | 37.8 (9.3) | 1.7 (0.5) | 4.5 (1) | 8.9 (1.8) | 8.9 (1.8) |
|  | Mineral (n=2) | 14.8 (5.4) | 1.5 (0) | 0.1 (0) | 3.9 (0) | 3.7 (1) | 3.7 (1) |
| Dry soil | O  (n=7) | 14.9 (5.1) | 20.5 (13.5) | 1.3 (0.9) | 6.6 (1.4) | 8 (2.6) | 5.5 (2.2) |
|  | Mineral (n=3) | 13.4 (6.3) | 1.8 (1.1) | 0.1 (0) | 5.1 (1.2) | 4.5 (1.8) | 4.5 (1.8) |
| Wet soil | O  (n=7) | 21.8 (4.8) | 35.2 (11.5) | 1.6 (0.5) | 5.1 (1) | 9.2 (1.6) | 8.1 (1.7) |
|  | Mineral (n=2) | 15.1 (4.6) | 1.6 (0.1) | 0.1 (0) | 3.7 (0.3) | 2.7 (2.4) | 2.7 (2.4) |
|  | ***Pinus banksiana*** | | | | | | |
| Control | O  (n=6) | 24 (4) | 23.1 (9.6) | 0.9 (0.5) | 4.1 (0.4) | 5.4 (0.5) | 5.4 (0.5) |
|  | Mineral (n=6) | 16.3 (4.7) | 0.8 (0.2) | 0 | 4.1 (0.7) | 4.6 (0.5) | 4.6 (0.5) |
| Dry soil | O  (n=6) | 12.5 (3.9) | 12.2 (9.8) | 0.9 (0.6) | 7.4 (0.3) | 5.5 (0.8) | 3.5 (0.8) |
|  | Mineral (n=6) | 14.4 (2.6) | 1.2 (0.3) | 0 | 4.6 (0.5) | 4.5 (0.8) | 4.5 (0.8) |
| Wet soil | O  (n=6) | 22.7 (0.5) | 29.8 (7.7) | 1.3 (0.3) | 4.3 (0.6) | 4.7 (0.8) | 3.5 (0.9) |
|  | Mineral (n=6) | 15.7 (3.7) | 1.2 (0.3) | 0 | 4.1 (0.7) | 5.2 (0.8) | 5.2 (0.8) |
|  | ***Populus tremuloides*** | | | | | | |
| Control | O  (n=6) | 14.8 (2.1) | 19.6 (9.7) | 1.3 (0.8) | 4.7 (0.3) | 7.3 (3) | 7.3 (3) |
|  | Mineral (n=4) | 12.5 (2.4) | 1.3 (0.3) | 0.1 (0) | 4.7 (1.1) | 5.3 (1.1) | 5.3 (1.1) |
| Dry soil | O  (n=6) | 10 (1.9) | 13.7 (9.3) | 1.4 (1) | 7.5 (0.5) | 7.5 (2.5) | 5.3 (2) |
|  | Mineral (n=4) | 9 (3.7) | 3.1 (2.5) | 0.5 (0.6) | 4.7 (1.1) | 3.5 (2.6) | 3.5 (2.6) |
| Wet soil | O  (n=6) | 14.7 (2.4) | 23.6 (15.2) | 1.6 (1) | 5.1 (0.2) | 6.9 (1.7) | 5.9 (1.9) |
|  | Mineral (n=4) | 15.2 (2.4) | 2.4 (1) | 0.1 (0) | 4.2 (0.3) | 3.8 (1.1) | 3.8 (1.1) |

| Table S5. Mean (SD) 2-pool decay coefficients for modelling microbial respiration post-burn. *M_1_* is the fractional active (or fast) C pool, and *k1* is the respiration rate constants for the fast C pool. | | | | | | | | | |
| --- | --- | --- | --- | --- | --- | --- | --- | --- | --- |
|  | Fast growth incubation | | | Affinity for post-fire environment incubation - Autoclaved | | | Affinity for post-fire environment incubation – Not autoclaved | | |
|  | k1 | M1 | | k1 | M1 | | k1 | M1 | |
| Unburned soil | 0.0036 (0.015) | | 0.096 (0.10) | 0.045 (0.039) | | 0.095 (0.061) | 0.014 (0.014) | | 0.28  (0.17) |
| Wet soil | 0.16  (0.12) | | 0.019 (0.013) | 0.046 (0.023) | | 0.096 (0.058) | 0.069  (0.12) | | 0.14  (0.13) |
| Dry soil | 0.016 (0.075) | | 0.034 (0.021) | 0.062 (0.027) | | 0.072 (0.039) | 0.058 (0.029) | | 0.074 (0.045) |

| Table S6. Mean (SD) 2-pool decay coefficients for modelling microbial respiration post-burn. *M_2_* is the fractional slow C pool, and *k2* is the respiration rate constant for the slow C pool. | | | | | | | | | |
| --- | --- | --- | --- | --- | --- | --- | --- | --- | --- |
|  | Fast growth incubation | | | Affinity for post-fire environment incubation - Autoclaved | | | Affinity for post-fire environment incubation – Not autoclaved | | |
|  | k2 | M2 | | k2 | M2 | | k2 | M2 | |
| Unburned soil | 0.0025 (0.00098) | | 0.89 (0.0025) | 0.00088 (0.00057) | | 0.89 (0.063) | 0.00039 (0.00063) | | 0.71  (0.17) |
| Wet soil | 0.0017 (0.00085) | | 0.97 (0.018) | 0.00088 (0.00054) | | 0.87  (0.06) | 0.00091 (0.00092) | | 0.84 (0.00091) |
| Dry soil | 0.00098 (0.00072) | | 0.05 (0.024) | 0.00053 (0.00025) | | 0.90 (0.064) | 0.00052 (0.00031) | | 0.89 (0.093) |

| Table S7. PERMANOVA results for full model for 16S for cDNA 24h after burning. | | | | | | | |
| --- | --- | --- | --- | --- | --- | --- | --- |
|  | Df | Sums of sqs | Mean sqs | F.Model | R^2^ | Pr(>F) | R^2^ single-component model |
| Dominant Vegetation | 2 | 0.000322 | 0.000161 | 2.0359 | 0.03604 | 0.001 | 0.03804 |
| Pre-burn horizon thickness (cm) | 1 | 0.000214 | 0.000214 | 2.6995 | 0.02389 | 0.002 | 0.02259 |
| pH | 1 | 0.000789 | 0.000788 | 9.9644 | 0.0882 | 0.001 | 0.13376 |
| Total C (%) | 1 | 0.000281 | 0.000281 | 3.5526 | 0.03145 | 0.001 | 0.0315 |
| Total N (%) | 1 | 0.00018 | 0.00018 | 2.277 | 2.277 | 0.003 | 0.03408 |
| Soil texture | 4 | 0.000593 | 0.000148 | 1.8746 | 0.06637 | 0.001 | 0.08829 |
| Burn treatment | 2 | 0.000222 | 0.000111 | 1.3994 | 1.3994 | 0.05 | 0.07969 |
| Soil horizon | 1 | 0.69043 | 0.000167 | 2.1112 | 0.01869 | 0.011 | 0.04257 |
| Residuals | 78 | 0.006172 | 7.91E-05 |  | 0.69043 |  |  |
| Total | 91 | 0.00894 |  |  | 1 |  |  |

| Table S8. PERMANOVA results for full model for 16S gDNA 24h after burning | | | | | | | |
| --- | --- | --- | --- | --- | --- | --- | --- |
|  | Df | Sums of sqs | Mean sqs | F.Model | R^2^ | Pr(>F) | R^2^ single-componentmodel |
| Dominant Vegetation | 2 | 0.001197 | 0.000598 | 3.017 | 0.05246 | 0.001 | 0.04618 |
| Pre-burn horizon thickness (cm) | 1 | 0.000746 | 0.000746 | 3.7606 | 0.03269 | 0.001 | 0.04093 |
| pH | 1 | 0.001238 | 0.001238 | 6.2422 | 0.05427 | 0.001 | 0.04924 |
| Total C (%) | 1 | 0.000848 | 0.000848 | 4.2757 | 0.03717 | 0.001 | 0.05109 |
| Total N (%) | 1 | 0.00035 | 0.00035 | 1.7636 | 0.01533 | 0.032 | 0.05114 |
| Soil texture | 4 | 0.002038 | 0.000509 | 2.568 | 0.0893 | 0.001 | 0.11346 |
| Burn treatment | 2 | 0.000666 | 0.000333 | 1.6781 | 0.02918 | 0.017 | 0.04097 |
| Soil horizon | 1 | 0.00046 | 0.00046 | 2.3204 | 0.02017 | 0.002 | 0.04873 |
| Residuals | 77 | 0.015275 | 0.000198 |  | 0.66942 |  |  |
| Total | 90 | 0.022818 |  |  | 1 |  |  |

:

| Table S9. PERMANOVA results for full model for microbial community composition at end of 5-week fast-growth incubation. | | | | | | | |
| --- | --- | --- | --- | --- | --- | --- | --- |
|  | Df | Sums Of Sqs | Mean Sqs | F.Model | R^2^ | Pr(>F) | Individual model (R^2^) |
| Dominant Vegetation | 2 | 0.000578 | 0.000289 | 2.0566 | 0.03493 | 0.001 | 0.03434 |
| Pre-burn horizon thickness (cm) | 1 | 0.000372 | 0.000372 | 2.6458 | 0.02247 | 0.002 | 0.02668 |
| pH | 1 | 0.001791 | 0.001791 | 12.7459 | 0.10823 | 0.001 | 0.11184 |
| Total C (%) | 1 | 0.000657 | 0.000657 | 4.6747 | 0.03969 | 0.001 | 0.04664 |
| Total N (%) | 1 | 0.000206 | 0.000206 | 1.4633 | 0.01243 | 0.071 | 0.04832 |
| Soil texture | 4 | 0.001023 | 0.000256 | 1.8206 | 0.06184 | 0.001 | 0.08092 |
| Burn treatment | 2 | 0.000693 | 0.000346 | 2.4648 | 0.04186 | 0.001 | 0.09525 |
| Soil horizon | 1 | 0.000269 | 0.000269 | 1.9115 | 0.01623 | 0.011 | 0.05924 |
| Residuals | 78 | 0.010957 | 0.00014 |  | 0.66233 |  |  |
| Total | 91 | 0.016543 |  |  | 1 |  |  |

| Table S10. PERMANOVA results for full model for microbial community composition at end of 6-month post-fire affinity incubation. | | | | | | | |
| --- | --- | --- | --- | --- | --- | --- | --- |
|  | Df | Sums Of Sqs | Mean | Sqs F.Mode | l R^2^ | Pr(>F) | Individual model (R^2^) |
| Dominant Vegetation | 2 | 0.001674 | 0.000837 | 3.845 | 0.03293 | 0.001 | 0.03286 |
| Pre-burn horizon thickness (cm) | 1 | 0.00088 | 0.00088 | 4.0411 | 0.0173 | 0.001 | 0.02287 |
| pH | 1 | 0.003791 | 0.003791 | 17.4128 | 0.07456 | 0.001 | 0.08042 |
| Total C (%) | 1 | 0.001445 | 0.001445 | 6.6355 | 0.02841 | 0.001 | 0.03597 |
| Total N (%) | 1 | 0.000711 | 0.000711 | 3.2656 | 0.01398 | 0.001 | 0.03738 |
| Soil texture | 4 | 0.002902 | 0.000726 | 3.3324 | 0.05707 | 0.001 | 0.07111 |
| Burn treatment | 2 | 0.000806 | 0.000403 | 1.8507 | 0.01585 | 0.001 | 0.04714 |
| Soil horizon | 1 | 0.000689 | 0.000689 | 3.1666 | 0.01356 | 0.001 | 0.05143 |
| Autoclave | 1 | 0.000938 | 0.000938 | 4.3073 | 0.01844 | 0.001 | 0.01831 |
| Residuals | 170 | 0.03701 | 0.000218 |  | 0.7279 |  |  |
| Total | 184 | 0.050845 |  |  | 1 |  |  |

Table S11. All responding OTUs. See all-responding-OTUs.csv

**Materials and methods**

Study region details: Mean temperatures range from 13 °C in the summer to -17.5 °C in the winter, and annual precipitation is 300-400 mm [1]. Most precipitation falls in the summer (June-September) months. Precipitation during the summer averages 50 mm per month but can range from as little as no precipitation during droughts to >140 mm per month [2]. For example, in Hay River, located approximately 35 km north of WBNP, summer precipitation in 2014 dropped to 27 mm per month from a summer monthly average of 41 mm and in September 2014, no precipitation was recorded. In Fort Smith, located on the eastern edge of WBNP, monthly precipitation < 3 mm was recorded in June 2007, August 2009, July 2010, and September 2011 [2].

Site details: Sites were located between 0.1 and 1 km from roads and > 0.5 km from other sampling sites. We used a Garmin GPSMAP 64 GPS finder to reach each designated location. Upon arriving, we confirmed the dominant tree species and recorded slope and aspect. Samples were collected across a 2 x 2-meter grid. A collapsible PVC pipe square was used to map out the sampling grid. After sampling most field sites, we produced a second selection of random points in a more limited region with field-validated tree species dominance. This second random sample was designed to address identified gaps in species dominance in the initial sample that were the result of errors and limitations in the map products used, resulting in a total of 19 sites, with 6-7 sites under each dominant vegetation type. At each site, ten soil cores (15.24 cm x 7.62 cm dia.) were collected using a soil core sampler with clear plastic core liners and plastic end caps (Product IDs 405.09 and 418.09; AMS, American Falls, ID, USA) every 1 m within (and two at the center of) a 2 m x 2 m grid.

Fire simulation: This heat flux was chosen to represent typical fire front propagation [3] and simulate a mid- to high-range crown fire (Thompson et al. 2015; Frankman et al. 2012; Butler et al. 2004). All the dry soil cores experienced some degree of combustion. Some but not all the wet soil cores ignited during the burn treatment.

pH: Briefly, we mixed mineral and organic soil with a 0.1 M CaCl_2_ solution in 1:1 and 1:5 soil:solution ratios, respectively. Samples were incubated at room temperature and vortexed every 20 minutes for 1 hour. Samples were centrifuged at 10,000 x g for 1 minute, and pH of the supernatant was measured using a pH electrode (Orion Star A215, Thermo Fisher Scientific, MA, USA) [7].

Total C and N: Medium and low organic content soil and clay (Product IDs B2178, B2152, and B2184; EA Consumables, Pennsauken, NJ, USA) were used to calibrate the instrument (Flash EA 1112 CN Automatic Elemental Analyzer (Thermo Finnigan, Milan, Italy)). We excluded C and N values from site 10 control, wet burn soil, and dry burn soil for all relevant analyses due to an error at time of measurement.

KOH base trap: Base trap absorption of grams of CO_2_ was calculated as the proportion (*P*) of the total base trap capacity for CO_2_ absorption (*CCA*) that is used, multiplied by the trap capacity (modified from Strotmann et al. 2004). *P* is calculated as (Equation 2):

P = (EC_init_ – EC_sample_)/ (EC_init_ – EC_sat_)	(2)

Where *EC_init_,* *EC_sample_*, *EC_sat_* are the electrical conductivity of the absorbing OH ion solution initially (0.5 M KOH), at the time of measurement, and at full saturation (0.25 M KOH), respectively. The base trap capacity for absorption of g CO_2_ is calculated as (Equation 3):

CCA = V x C x R x M	(3)

Where *V* is the volume of the absorbing OH ion solution (0.015 L), *C* is the initial concentration of KOH, *R* is the mole to mole ratio of KOH present to CO_2_ absorbed (0.5), and *M* is the molecular mass of CO_2_ [8].

Incubation setup: For both incubations, soil was packed to match the wet bulk density of the original core as recorded during the destructive sampling process post-burn treatment. Bulk densities ranged from 0.005-0.29 g cm^-3^ for the O horizon and 0.25-0.61 g cm^-3^ for the mineral soil. Finally, deionized water was added to each sample to bring all samples to equivalent soil moisture across all incubations from a given site. Target soil moistures were based on a water holding capacity of 175% g H_2_O per g dry soil for O horizons and 55% water filled pore space for mineral soil. Seven of the 19 sites contained O horizons that we classified as Fibric Histosols, in which case we used a target moisture of 285% g H_2_O per g dry soil, which is consistent with peatland moistures [9].

RNA and DNA: RNA and genomic DNA (gDNA) co-extractions were performed for each soil horizon of each sample with a blank extraction (identical methods but with empty tubes) for every 8 samples using RNeasy PowerSoil Total RNA Kits and RNeasy PowerSoil DNA Elution Kits (QIAGEN, Germantown, MD, USA), respectively. In brief, 0.5 – 1 g of O horizon or 2 g of mineral soil was added to a PowerBead tube. A phenol/chloroform nucleic acid extraction was performed following manufacturer’s instructions. Homogenization and lysis were done using a FastPrep-24™ 5G bead beating grinder and lysis system (MP Biomedicals, Irvine, CA, USA) at a frequency of 6.5 m s^-1^ for 45 s at room temperature.

Residual gDNA contamination was removed from the RNA extracts using DNase Max Kits (QIAGEN, Germantown, MD, USA) following manufacturer’s instructions. In brief, 40 µL of sample was combined with a DNase enzyme, buffer, and nuclease-free water and incubated at 37 °C for 20 min. The sample was then combined with 5 µL of DNase removal resin, incubated for 10 mins on a vortex adapter set to low, and centrifuged at 13,000 x g for 1 min. Supernatant was collected in a clean tube for further analysis.

RNA reverse transcription was carried out using Invitrogen SuperScript IV VILO Master Mix (ThermoFisher Scientific, Waltham, MA, USA) following manufacturer’s instructions. A control reaction using no reverse transcriptase enzyme was carried out on one of every three RNA extracts to ensure minimum contamination by gDNA in the RNA sample. A Quant-iT RiboGreen RNA Assay Kit (ThermoFisher Scientific, Waltham, MA, USA) was used to assess copy DNA (cDNA) concentration and gDNA contamination levels.

PCR: In brief, PCR mixes contained 1.25 µL 515f forward primer (10 µM), 1.25 µL 806r reverse primer (10 µM), 1 µL cDNA or gDNA template, 12.5 µL Q5 Hot Start High-Fidelity 2X Master mix (New England BioLabs INC., Ipswich, MA, USA), 1.25 µL Bovine Serum Albumin (BSA) (20 mg mL^-1^) (VWR, Radnor, PA, USA), and 7.75 µL nuclease-free water, in a 96-well plate. The plate was sealed and placed on an Eppendorf Mastercycler nexus gradient thermal cycler (Hamburg, Germany). Reactions were run at 98 °C for 2 minutes + (98 °C for 30 seconds + 58 °C for 15 seconds + 72 °C for 10 seconds) x 30 cycles + 72 °C for 2 minutes and 4 °C hold. The quality of PCR amplicon triplicates was assessed using gel electrophoresis, and triplicates were then pooled, purified, and normalized using SequalPrep Normalization Plate Kits (96-well) (ThermoFisher Scientific, Waltham, MA, USA). Samples were then pooled, and library cleanup was performed using a Wizard SV Gel and PCR Clean-Up System (Promega, Madison, WI, USA). The pooled library, including blanks, was submitted to the UW-Madison Biotechnology Center (Madison, WI, USA) for 2x250 paired end Illumina MiSeq sequencing. The library was sequenced twice using identical protocols to improve sequencing depth. Reads from the two sequencing runs were pooled by sample after sequence processing and before analysis.

Sequence processing: We quality filtered, trimmed (left trim 13 for both reads; truncation length 182 for forward reads, 161 for reverse reads), and dereplicated, learned errors (1M reads, randomized), picked operational taxonomic units (OTUs), and removed chimeras (consensus method) using dada2 [10] as implemented in Quantitative Insights Into Microbial Ecology (QIIME2) [11]. A total of 18,830,558 reads were obtained from amplicon sequencing. 6,360,490 reads were removed during the filtering processing, and taxonomy was assigned to the remaining 12,470,068 reads (24,940 mean 16S reads per sample) using the naïve Bayes classifier [12] in QIIME2 with the aligned 515f-806r region of the 99% OTUs from the SILVA database (SIVLA 138 SSU) [13–15]. We excluded 1 cDNA and 2 gDNA samples from further analysis due to low 16S reads per sample (<1000).

Statistics: We compared community composition across samples using weighted UniFrac dissimilarities on RNA- and DNA-based relative abundances and tested for significant effects of horizon (O horizon versus mineral soil), dominant vegetation, pre-burn horizon thickness, pH, total C and N, soil texture, and degree hours (DH) using a permutational multivariate ANOVA (PERMANOVA using the *adonis* function in *vegan* [16]) and used the *betadisp()* function to test if variables differed in their dispersion. We used single-component models to compare the R^2^ for each factor.

We used the rrnDB RDP Classifier tool (version 2.12) to obtain a mean 16S rRNA gene copy number for each genus in the fast growth dataset [17]. Briefly, for taxa with and without a genus-level assignment, we used the genus mean copy number or the mean copy number for all other taxa in this study, respectively, in the RDP database as the predicted copy number. We normalized OTU counts by dividing by the predicted copy number and summed the product of the predicted copy number and each OTU relative abundance for all OTUs in each sample to calculate the community weighted mean rRNA gene copy number.

**References**

1.	ESWG. A national ecological framework for Canada. *Agriculture and Agri-Food Canada, Research Branch, Centre for Land and Biological Resources Research and Environment Canada, State of the Environment Directorate, Ecozone Analysis Branch*. 1995. Ottawa, Ontario/Hull, Quebec, Canada.
